# DNA tension-modulated translocation and loop extrusion by SMC complexes revealed by molecular dynamics simulations

**DOI:** 10.1101/2021.03.15.435506

**Authors:** Stefanos K. Nomidis, Enrico Carlon, Stephan Gruber, John F. Marko

## Abstract

Structural Maintenance of Chromosomes (SMC) complexes play essential roles in genome organization across all domains of life. To determine how the activities of these large (≈50 nm) complexes are controlled by ATP binding and hydrolysis, we developed a molecular dynamics model that accounts for conformational motions of the SMC and DNA. The model combines DNA loop capture with an ATP-induced “power stroke” to translocate the SMC complex along DNA. This process is sensitive to DNA tension: at low tension (0.1 pN), the model makes loop-capture steps of average 60 nm and up to 200 nm along DNA (larger than the complex itself), while at higher tension, a distinct inchworm-like translocation mode appears. By tethering DNA to an experimentally-observed additional binding site (“safety belt”), the model SMC complex can perform loop extrusion (LE). The dependence of LE on DNA tension is distinct for fixed DNA tension vs. when fixed DNA end points: LE reversal occurs above 0.5 pN for fixed tension, while LE stalling without reversal occurs at about 2 pN for fixed end points. Our model matches recent experimental results for condensin and cohesin, and makes testable predictions for how specific structural variations affect SMC function.

## INTRODUCTION

Chromosomes in all living cells contain tremendously-long DNA molecules, ranging in size from megabases (millimeters) in bacteria and other microbes, to gigabases (meters) in some animals and plants. Many lines of evidence have long pointed to DNA or chromatin loop formation as a fundamental organizing principle of chromosome folding, and it has now become clear that the Structural Maintenance of Chromosomes (SMC) protein complexes are key drivers of DNA looping (1, 2, 3). SMC complexes (SMCCs) are found in eukaryotes, bacteria and archaea, and possess a distinctive ring-like architecture. The rings include two SMC proteins with long, flexible regions that connect dimerization “hinge” domains at one end to an ATP-binding domain “head” at the other end (Fig. 1A).

**Figure 1.**
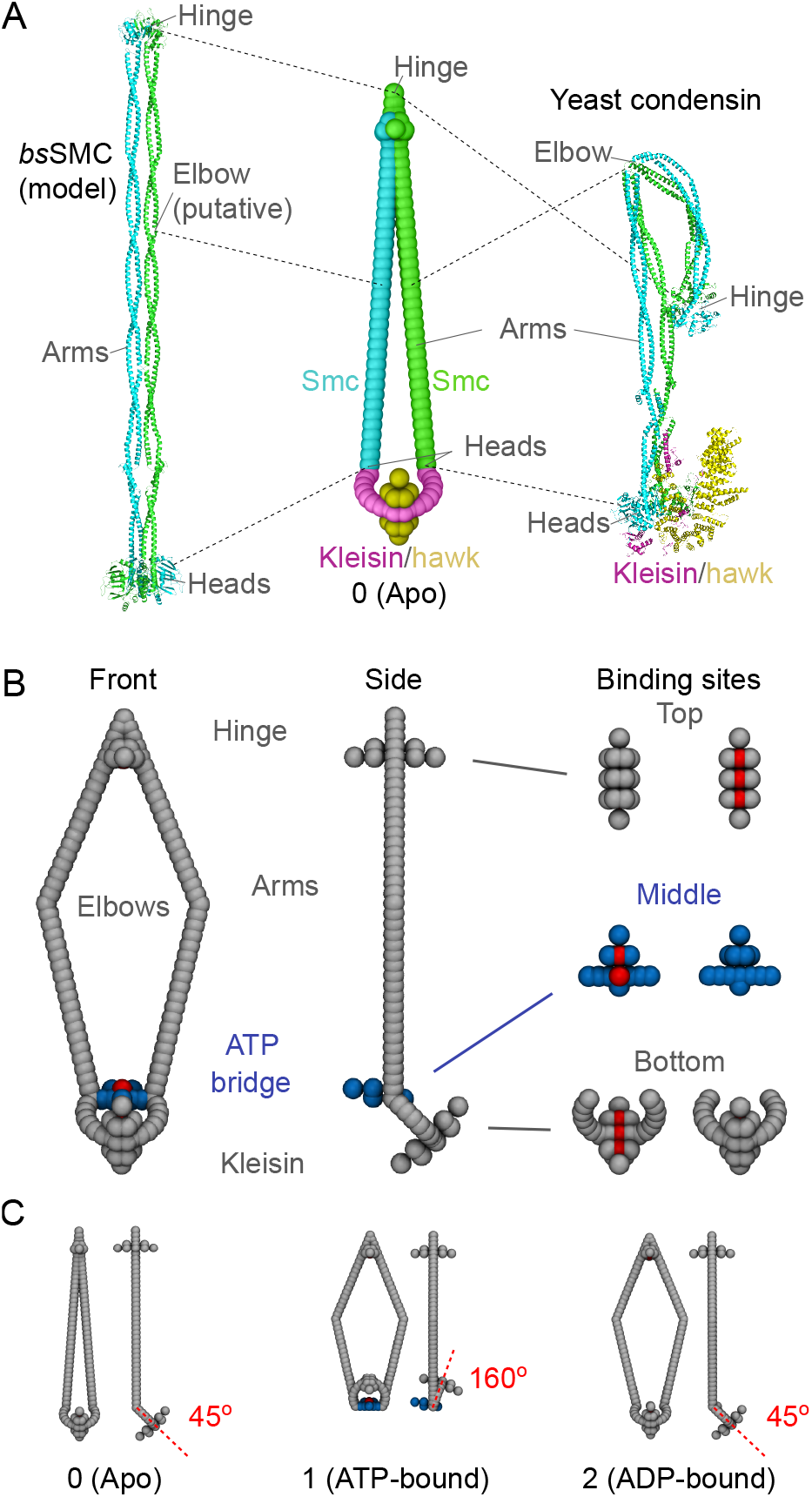
(A) SMCC structure in the ATP-unbound “apo” or closed state. Left: Composite structural model of bsSMC (19), showing the two SMC proteins in cyan and green, the dimerization “hinge” domain (top, PDB structure 4RSJ), the long coiled-coiled arms (PDB 5XG2 and 5NNV) with putative “elbow” domain, and ATP-binding/ATPase “heads” (PDB 5XEI). Additional subunits are shown in purple. The two SMC proteins are adjacent, leaving little space between them. Right: Structural model of budding yeast condensin based on cryo-EM data (PDB 6YVU) (20) following the same coloring scheme, with the kleisin in purple and the additional “hawk” subunit shown in yellow; for yeast condensin the coiled-coils are bent in the apo configuration, with the hinge domain near the lower part of the arms. Center: Simulation model of SMCC in the closed state, with SMC protein arms near one another. (B) Geometry of the simulation model, with coiled-coil arms “open”. A front and side view are shown on the left and in the middle, respectively. Beads that are repulsive and attractive to DNA are shown in light gray and red, respectively, while the DNA-repulsive ATP-bridge beads are indicated in blue. The “hinge” domain, the ATPase-bridge domain, and the kleisin domain are shown separately on the right, with top and bottom views of the DNA-binding domains shown. DNA-binding sites are shown in red (bottom of “hinge” domain, top of “ATPase bridge” domain, top of domain). Note that, this open SMCC configuration (open arms, ATPases bridged *and* kleisin at a 45-degree angle) is not an actual state of the model, but is used for illustration purposes. (C) The three actual structural states of the SMCC, for no ATP bound or “apo” (left), ATP bound “open” state (center) with folded kleisin, and ADP-bound “ATPase bridge open” state (right). In the apo state, the top, middle and bridge DNA binding sites are off; in the ATP-bound state the top, middle and bridge sites turn on; and in the ADP-bound state the middle, bridge and lower sites turn off.

SMCCs undergo a series of conformational changes driven by ATP hydrolysis while interacting with DNA, leading to physical organization and folding of chromosomes. Cohesin and condensin are the two most studied eukaryotic SMCCs: condensin (4, 5) folds and compacts chromosomes via formation of tightly-packed tandem loops (6, 7, 8), while cohesin (9) holds together identical copies of DNA formed during mitosis (10, 11) and was also found to play an important role in gene expression (12, 13). The bacterium *B. subtilis* posesses the bsSMC complex (14, 15), and *E. coli* has the complex MukBEF (16, 17, 18). These bacterial SMCCs are structurally similar to condensin and cohesin, and fold chromosomes into highly-compacted, disentangled structures.

SMCCs are large (≈50 nm) and conformationally flexible. They are assembled as dimers of two SMC proteins which each have a dimerization domain at one end and an ATP-binding and hydrolysing domain at the other. The binding of the two SMC proteins at their dimerization domains forms the “hinge” of the SMCC, while at the ATPase domain the SMCC has a more complex structure capable of large conformational changes. Figure 1A shows structural models for bsSMC and yeast condensin (19, 20) along with our MD model, all for the “closed” configuration with SMC arms adjacent to one another (Table S1 of Supplementary Data lists corresponding structural features of these two SMCCs). This configuration corresponds to the “apo” state where no ATP is bound.

A characteristic of all SMCCs is the presence of a kleisin subunit, which links the two SMC proteins to form a tripartite ring (21). *In vivo* experiments have established that DNA can be threaded through this protein ring (22, 23, 24). When ATP is bound by the ATPase domains of each of the two SMC proteins, the two heads can bind together to form a bridge, transforming the single ring of the SMCC to a two-compartment ring, consisting of a large “SMC” compartment and a smaller “kleisin” compartment (Fig. 1B). The kleisin compartment is known to be able to trap DNA (25). ATP hydrolysis unlinks the ATPases, returning the ring to a single-compartment conformation, and then finally b ack t o a conformation where the coiled-coil arms are adjacent to one another. Thus, the ATP binding, hydrolysis and release cycle is coupled to large-scale conformational changes of the SMCC.

A structural feature that varies among different SMCC species is the degree of folding of the coiled-coils at the elbow sites, to bring the hinge domain towards the kleisin region (*e.g.*, Fig. 1A, yeast condensin). Yeast condensin (20), cohesin (26, 27) and *E. coli* MukBEF (27, 28) are known to have this folding, while SMC5/6 (29) and *B. subtilis* bsSMC (30) are thought not to have it. As will be discussed, our model is not strongly dependent on the precise geometry of the complex, other than having the coiled-coils adjacent and the upper compartment “closed” in the apo state, and having the upper compartment open when ATP is bound.

It has been hypothesized (1, 2, 6, 7, 12, 13) that SMCCs are able to translocate along the DNA double helix (or along chromatin) in the manner of a molecular motor. Such a motor can perform active loop extrusion, *e.g.*, by simply binding to one spot on the DNA while translocating (thought to be the case for yeast condensin (31)). Indeed, a series of experiments have observed ATP-dependent DNA compaction by condensin (32, 33, 34), translocation by condensin (35), and loop extrusion by condensin (36) and cohesin (37, 38, 39, 40). The mechanism by which SMCCs perform these functions is unclear, although it must be based on ATP binding and hydrolysis, coupled to protein conformational change, and in turn coupled to DNA conformational fluctuation, which combine to produce translocation and loop extrusion.

Here, we present a coarse-grained molecular dynamics (MD) model which provides a generic description of SMCC activities. It is based on prior analytical work on SMCC translocation and loop extrusion (41), but is able to take into account aspects of SMCC-DNA interactions which are difficult or impossible to deal with in a purely analytical framework, notably the synergy between conformational flexibilities o f the S MC complex and t he D NA it is moving along. The MD model is constructed from known features of SMCC-DNA interactions and the SMCC ATPase cycle, and contains enough detail to make a wide range of tests and predictions of SMCC behavior (Fig. 1A-B). A number of models of SMCC function were proposed prior to our segment-capture model (41), based on mechanisms including “inchworm” motions (42), a rotary step of a SMC subunit (43), and a non-motor based loop-ratchet (44). However, these works did not consider how the ATP-cycle-coupled conformational change of SMCCs and DNA random fluctuation work together to lead to active translocation and loop extrusion as observed experimentally, which is the focus of this paper.

The basic DNA-segment-capture mechanism (41) qualitatively explains existing experiments on SMCC translocation along DNA, and subsequent theoretical work incorporating similar mechanisms involving binding at one DNA site with capture of a second, distant DNA (45, 46) has led to concordant results. An alternative ‘scrunching’ model has also been proposed, based on the idea that DNA might be handed over from the hinge to the heads (or vice versa) via folding of the SMCC at an “elbow” joint (35, 47, 48, 49). However, elbow folding has not been observed for Smc-ScpAB and Smc5/6 (30, 50, 51). The positions of the elbow in condensin, cohesin and MukBEF are different, with hinge meeting arms, joints and heads, respectively, in those complexes, and the structure of the elbow is not conserved. This suggests elbow bending being a feature arising from divergent evolution, rather than being a feature fundamental to translocation and loop extrusion (20, 27, 52). We also note that scrunching and other SMCC-walking-based models (49) have difficulty generating steps along DNA that are larger than the complex itself, as has been observed for SMCCs (53); in contrast steps larger than the SMCC itself occur naturally in models that are based on DNA loop capture (41, 45).

Here, we report simulations of a MD realization of our analytical model that reveal new phenomena. We find that the flexibility of the SMC coiled-coils allows an alternative “inchworm” mode of translocation to come into play, when the underlying DNA is under too much tension to permit DNA segment bending to easily occur (roughly for DNA tension above 1 pN). Anchoring of the DNA to the SMCC during translocation gives rise to loop extrusion, and we find that existing experiments observing loop extrusion on DNAs anchored at two fixed points (thus at *fixed DNA extension*) are also readily described, with loop extrusion persisting up to tensions of about 2 pN. Remarkably, we also find that, if a similar experiment is carried out at *fixed DNA tension*, different mechanoenzyme behavior is observed, with true stalling followed by “loop de-extrusion”, as force is increased beyond about 0.5 pN. Thanks to the role played by their distinctive structure in SMCC function, there are a number of conceivable modifications to SMCCs which should change their mechanical properties. We explore a few of these using our MD framework, predicting results for experiments on varied or mutated SMCCs.

## MATERIALS AND METHODS

### Model Geometry

Our coarse-grained MD model is based on structural features common across SMCCs (19, 41). The model was designed to be a minimal model incorporating the key geometrical and topological elements necessary for translocation, namely broken spatial and time symmetries. We will show that the behavior of the model is robust to appreciable parameter variation, but the key elements (the broken symmetries, the enzyme conformational cycle, and the topological coupling of the enzyme to DNA) are essential. It consists of individual rigid bodies (Fig. 1B), which interact through bond, angle and dihedral potentials.

The geometrical details of the model have been chosen based on structural data for SMCCs as follows. Each of the two 50-nm-long coiled-coil arms consists of two straight segments, joined at a flexible “elbow” (27), which gives them the ability to open and close (19, 30). The two SMCs are connected at the top by a dimerization “hinge”. The hinge region of many SMCCs is known to contain a DNA-binding locus (54, 55, 56, 57, 58, 59, 60), although some studies indicate that having a DNA binding site at the hinge may not be essential (25). The SMCs can also be connected at their ATPase domain “heads”, via two bound ATPs, and by a kleisin subunit (19). The latter establishes the overall ring structure of the SMCC, and formation of the ATPase-ATP-ATPase “bridge” (shown in blue in Fig. 1B) can divide the ring into two compartments (25, 28, 61, 62).

We will refer to the DNA-binding sites of the hinge, ATPase bridge and kleisin as *top*, *middle* and *bottom* binding sites, respectively (Fig. 1B, right). We make special note of the middle site at the bottom of the upper compartment, which requires ATP binding and engagement. This is a highly-conserved feature across SMCCs, and has been found to be essential for translocation in bsSMC (25, 26, 62, 63, 64, 65). We use energetic binding to model SMCC-DNA interactions, but to some degree these sites, in particular the lower one, may sterically trap or embrace DNA (25, 62).

### SMCC Structural states

The model SMCC has three distinct structural states, corresponding to the ATP-unbound (apo), ATP-bound, and ATP-hydrolyzed/ADP-bound states of the ATPase (Fig. 1C) (41). In the *apo state* (0), the two arms and the upper compartment are closed, the ATP/ATPase bridge is open, the hinge DNA binding site is turned off and the lower compartment is folded by 45 degrees. In the *ATP-bound state* (1), which occurs upon ATP binding, the ATP bridge closes, the two arms open, and the lower compartment folds by an angle of 160 degrees. These conformational changes are supported by ultrastructural experimental data. Angular conformational changes of the lower compartment region of similar symmetry to those in our model and associated with ATP binding have been observed in cryo-EM studies of condensin and cohesin (20, 26, 28, 52, 65). In the case of yeast condensin this angular change involves the kleisin Brn1 and the Ycs4 HEAT-repeat subunit, and not the Ycg1 HEAT-repeat subunit (20, 66). Partial opening and closing of the coiled-coils of the long SMC proteins have also been observed (20, 66). Intriguingly, while high-speed AFM experiments have observed a “folding” transition of condensin where the hinge moves to contact the Walker ATPase heads (47) this has not been observed in the recent cryo-EM experiments.

The opening of the arms makes the top DNA-binding site at the dimerization hinge accessible, which we model by turning on the top binding site of the open upper compartment. The same transition also turns on the middle DNA-binding site, at the lower part of the compartment. Finally, in the *ADP-bound state* (2), the two arms remain open, the bottom and middle binding sites are turned off, the folding angle of the kleisin is reduced back to 45 degrees and the ATPase bridge opens, and no longer separates the two compartments. Note that, the folding of the lower compartment is asymmetric, which is necessary for the motion of the SMCC to be directional.

The transition from each SMCC state to the next is modeled through changes of the equilibrium angles, which lead to opening/closure of the arms and folding of the kleisin. The bridge addition/removal is, likewise, modeled by the addition/removal of excluded volume interaction terms in the bridge beads. The transition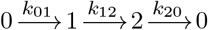 take place at rates *k*_01_, *k*_12_ and *k*_20_, respectively. In practice, the simulation is maintained in a state *i* for a time interval *t_i_*, with *t_i_* a random variable selected from an exponential distribution 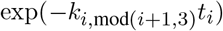. Within each state, the system evolved through ordinary MD with an implicit solvent Langevin thermostat. The configurations of Fig. 1(C) are the minimal-energy states of the SMCC. Due to interactions with DNA and with the thermal environment, the SMCC structure fluctuates during simulations (see Supplementary Data Movies S1, S2 and S3). Parameters used to define the geometries and DNA-binding site dynamics for the different SMCC states are in Table S2 and Fig. S1 of Supplementary Data.

### Translocation mechanism

In our model, the SMCC translocates along DNA via a segment-capture mechanism (41), as illustrated in Fig. 2A. DNA is threaded through the lower compartment and bound at the bottom binding site. This is the apo state, with the two arms closed and the ATPase bridge open. The passage of DNA through the tripartite ring ring has been established in topology-sensing experiments on a range of SMCCs (22, 23, 25, 28, 67).

**Figure 2.**
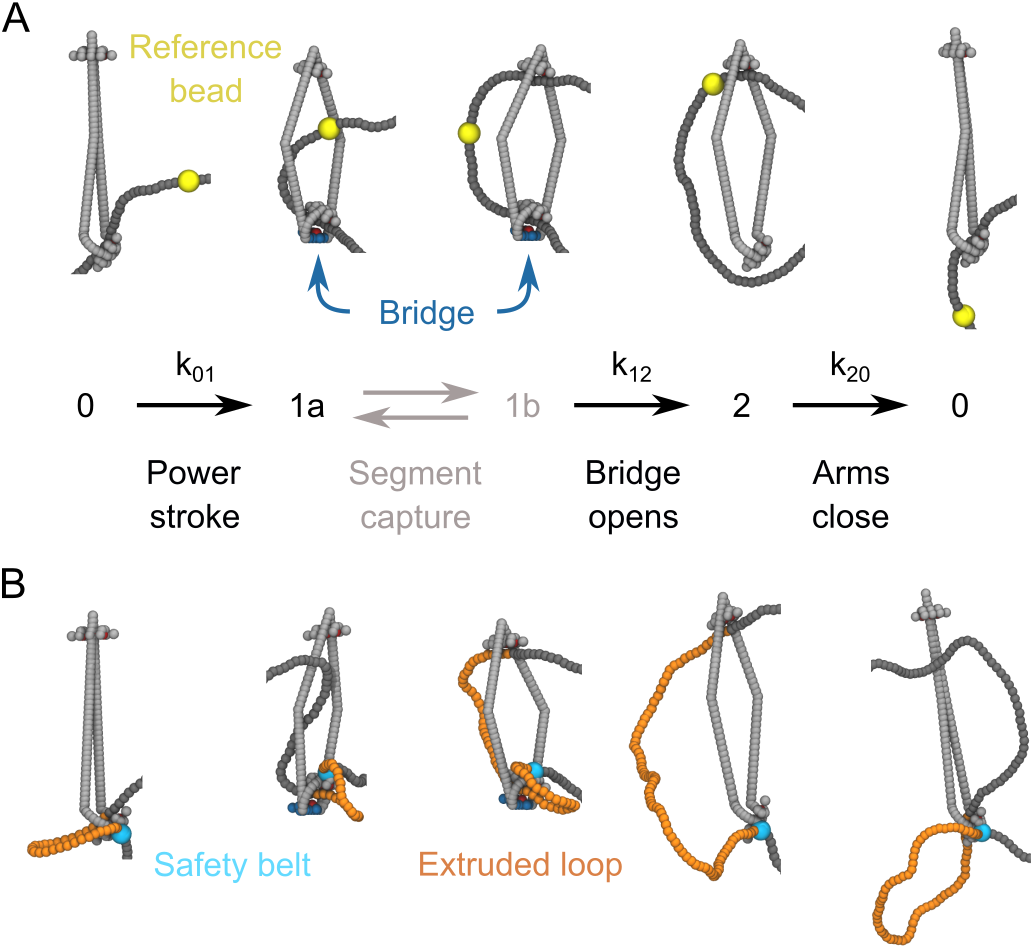
Sample MD simulation conformations of SMCC and DNA during (A) translocation and (B) loop extrusion. For translocation and loop extrusion the SMCC undergoes the same type of conformational changes, cycling through the three states labeled 0, 1 and 2 (Fig. 1C). In the transition from 0 to 1 the kleisin folds and the ATPase bridge forms, directing the DNA inside the SMCC upper compartment. In the transition to 2 the bridged ATPases separate and the DNA detaches from the now-nonexistent bottom binding site on the SMCC. The closure of the arms pushes the DNA back to the bottom of the SMCC at the end of the cycle. The net effect is the translocation of DNA, shown here by the displacement of a reference bead (yellow). In the simulations, backward steps can be observed, particularly at higher DNA tensions (see text). In the loop extrusion case (B) the interaction DNA/SMCC is as in (A), but one site of DNA is permanently attached to the exterior of the kleisin subunit (cyan bead indicates this “safety belt” binding site), hence the translocation along the DNA extrudes a loop (yellow).

Upon transitioning to the ATP-bound state, the lower compartment folds, bringing DNA close to the middle binding site, to which it can efficiently bind (state 1a). This plays the role of a power stroke, as it forcibly pushes a DNA segment into the upper compartment. Notably, a recent cryo-EM structure of MukBEF has revealed that DNA is bound to that SMCC in the manner shown in state 1a, with the DNA passing through the asymetrically bent lower compartment (28). Subsequently, conformational fluctuations can lead distal DNA to bind to the upper binding site (state 1b), thus “capturing” a bent DNA segment (one could describe this captured segment as a small loop, but we will avoid that to better distinguish it from the larger, extruded DNA loops).

Following segment capture upon ATP hydrolysis, the ATPase bridge opens, and conformational changes in the lower compartment lead to the middle and bottom binding sites being no longer active. Given that the DNA sequestration in the lower compartment may be due to steric trapping (25, 62), the release of DNA from the lower site could be a natural consequence of the opening of the bridge. With the bridge gone, the captured DNA segment can release bending stress by escaping through the bottom of the complex. Finally, the SMCC returns to its apo state, with the closing of the two arms entropically pushing the DNA back to the now-reactivated bottom binding site. This sequence of states, ending with transport of the distal end of the captured segment from the top to the bottom of the SMCC protein loop, results in translocation of the SMCC along the DNA (indicated by the yellow reference DNA bead, Fig. 2A).

An essential feature of this reaction cycle is that the DNA segment is never passed through the tripartite SMC-SMC-kleisin protein ring (this may occur during loading, but not during the translocation cycle). In addition to being supported by experimental observations of topological linkage of SMCCs to DNA mentioned above, this topological feature is key to the high degree of processivity and conserved translocation directionality of SMCCs along their left-right symmetric dsDNA substrate observed experimentally (35). Without this linkage, SMCCs would be unable to preserve their direction of translocation along dsDNA (41, 66). We also emphasize that in this model the SMCC transitions are decoupled from DNA motion, in the sense that steps in the protein conformational cycle are not contingent on *e.g.*, the presence of DNA bound at particular SMCC loci. At present there is no evidence for such “synchronizing” interactions although they could certainly be added to the model. The absence of this type of synchronization in the present model means that it is possible for the segment capture process to fail, *i.e.*, for futile SMCC cycles to occur.

### Loop-extrusion mechanism

Given translocation, loop extrusion may occur in a variety of ways (41, 68). In this paper, we assume that a DNA segment (cyan bead in Fig. 2B) is attached to the exterior of the kleisin subunit, as is thought to be accomplished by the “safety belt” of yeast condensin. This DNA binding site is formed by Ycg1 and part of Brn1 (31), distinct from the regions of the Brn1 and Ycs4 involved in the ATP-driven angular conformational change observed in cryo-EM experiments (20, 66). Combining safety-belt site binding with the threading of DNA through the lower compartment leads to formation of a DNA loop. In this situation, translocation, as described above, leads to asymmetric loop extrusion, as the unattached end of the loop translocates along DNA. We note that, establishment of this mode of loop extrusion requires passage of the DNA through the SMC-kleisin protein ring (*topological loading*), possibly involving opening of a SMC-kleisin “gate” (65, 69). We explore alternate loop-extrusion mechanisms compatible with our translocation model in the Discussion.

### Computational Methods

#### MD

All simulations were performed in LAMMPS (70), using a standard velocity-Verlet integration scheme coupled to a Langevin thermostat. The simulation temperature was 300 K, with a damping parameter of 0.5 ns, while the simulation timestep was 0.2 pLsec (we use the units of Lsec to refer to LAMMPS time to distinguish from real experimental time measured in seconds).

#### DNA

DNA was modeled as a semiflexible bead-and-spring polymer (dark gray beads in Fig. 2). Each bead represented 5 bp with a mass of 0.005 ag (3 kDa), while successive beads were connected through finitely-extensible springs, with a rest length of 1.7 nm We emphasize that our simulations results are numerically independent of the bead masses due to the fact that the SMCC and DNA dynamics of interest here are on a much longer time scale than the relaxation time of bead velocity. A demonstration of the independence of kinetics of the DNA on bead mass is included in Supplementary Figure S5. The DNA stiffness was modeled through angle interactions among three successive beads, yielding a persistence length of 50 nm. Excluded-volume interactions were taken into account by means of truncated and shifted Lennard-Jones interactions, with an interaction distance of 3.5 nm, *i.e.*, the effective DNA diameter at physiological salt conditions (71). A total of 301 beads were used, corresponding to a 1.5-kbp sequence with a contour length of 510 nm.

#### SMCC

The SMCC consisted of 7 rigid bodies (Fig. 1B): four coiled-coil segments, the top binding site, the ATP bridge together with the middle binding site, and the kleisin subunit together with the bottom binding site. These interacted with each other by means of bond, angle and dihedral potentials. Each bead had a diameter 1.5 nm, and the total mass of the SMCC was chosen to be 0.25 ag (150 kDa). The coiled-coil arms were made of two connected, 25-nm-long, straight segments, interacting through a harmonic angle potential of 30 *k*_B_*T* /rad^2^ stiffness. The upper binding site consisted of a total of 17 beads. Three of these were chosen to be attractive to DNA (red beads), while the rest of them were repulsive (light gray beads). This ensured that DNA could only bind from a single direction. The ATP bridge was made of 2 attractive (red) and 6 repulsive beads (blue) attached to a 7.5-nm-long, straight segment (blue). The kleisin subunit consisted of 3 attractive (red) and 14 repulsive beads (light gray) attached to a circular arc of radius 7 nm (light gray). The top binding site, the ATP bridge and the kleisin subunit were kept aligned by means of bond and dihedral interactions.

The attraction of DNA by the binding sites was modeled with a truncated and shifted Lennard-Jones interaction, with a potential depth of 3.2 *k*_B_*T* for each top- and middle-binding-site bead, and 11 *k*_B_*T* for the bottom-site beads. In order to control the angle between the ATP bridge and the lower half of each arm, harmonic angle potentials were used, with a stiffness of 100 *k*_B_*T*/rad^2^. The two arms were made bendable by introducing similar harmonic angle potentials, with a stiffness of 30 *k*_B_*T*/rad^2^. The asymmetric folding of the kleisin subunit was achieved through two harmonic dihedral interactions, with stiffness constants of 60 *k*_B_*T*/rad^2^ and 100 *k*_B_*T*/rad^2^. The SMCC cycled stochastically through the three different states, apo (0), ATP-bound (1) and ADP-bound (2), with mean dwell times of *τ*_01_ = 0.4 *μ*Lsec, *τ*_12_ = 1.6 *μ*Lsec and *τ*_20_ = 0.4 *μ*Lsec (LAMMPS time units). Instantaneous dwell times were drawn from exponential distributions.

For the determination of captured DNA loop sizes we identified where the upper SMCC compartment encircled DNA. In particular, we located the DNA bead closest to the center of mass of the upper compartment, and also the one closest to the top binding site, and compared the two distances. This determined whether the system was in state 1a or 1b, *i.e.*, DNA was closer to the center of mass or the top binding site, respectively. The loop end was associated with the smallest-proximity DNA bead.

## RESULTS

### Translocation

We performed MD simulations for translocation of an SMCC along DNA under varying DNA tension, *i.e.*, with stretching force applied to the two DNA ends (see Supplementary Data, Movies S1 and S2). Figure 3A shows time traces of SMCC positions along the DNA track for three different DNA tensions. The position is here defined as the coarse-grained DNA bead with minimal distance from the SMCC bottom site. This quantity changes slowly when the SMCC is in the states 0 and 1, while it shifts abruptly with the closing of the arms during the transition 2 → 0. At the smallest tension in Fig. 3A (0.1 pN) the steps shown are all forward, while an increasing number of backward steps are observed as the tension increases. The timescale shown in the figure is based on experimental data of Ref. (36) and is applied by rescaling our simulation timescale to have each cycle correspond to 0.13 sec (see below). Fig. 3B shows the observed translocation step size (blue points), averaged over many repeated cycles. As expected, the translocation slows down with increasing values of DNA tension, since it requires substantial bending of DNA (see Fig. 2A), which becomes progressively unfavorable as the latter gets stretched. Interestingly, the translocation of the SMCC does not halt at large DNA tensions, but instead reaches a plateau value of about 30 nm per cycle. This is due to the asymmetric folding of the lower compartment in the ATP-bound state, which may be viewed as a power stroke (state 1a in Fig. 2A), and is essentially unaffected by the physiologically-relevant forces (< 10 pN) considered here.

**Figure 3.**
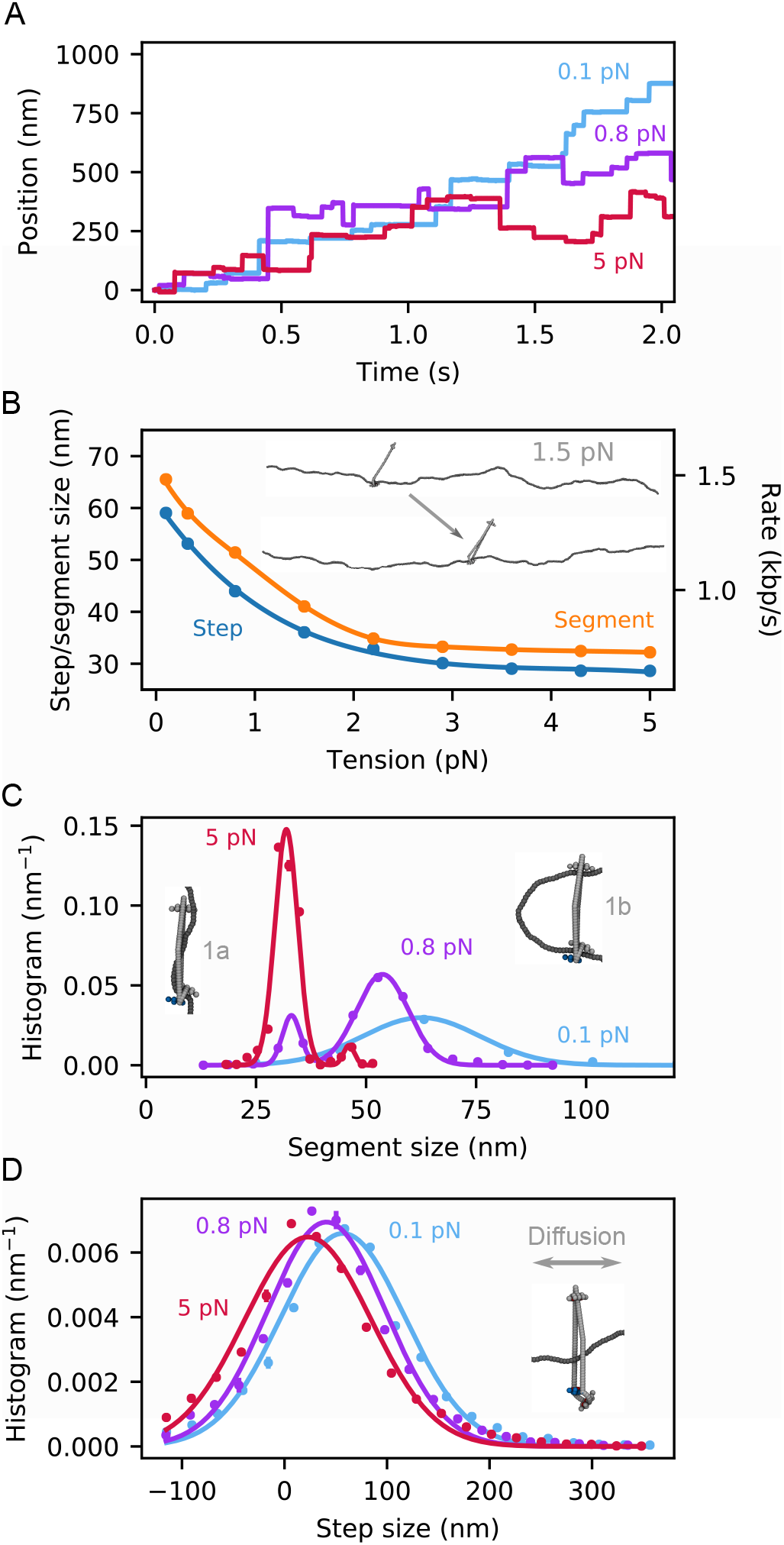
DNA translocation by SMCC. (A) Representative traces of the SMC position along the DNA track for three selected values of the DNA tension. These were obtained by stacking together independent SMC cycles under the same initial conditions. (B) Mean step size (blue points) and captured segment length (ATP-bound state 1 in Fig. 2A, shown with orange points) dependence on DNA tension. Solid lines are spline interpolations, while the right axis converts the mean step size to translocation rate, using the conversion factor derived in Fig. 4. The inset shows the before- and-after configurations for a typical cycle at 1.5 pN. (C) Segment-size distribution for some selected tensions (normalization is chosen so that the distribution integrates to 1). The solid lines are single- and double-Gaussian fits to the data; for the double-peak distribution the left and right peaks correspond to substates 1a and 1b respectively (inset snapshots; see text). (D) Step-size distribution for some selected tensions. Notice the difference in the length scale and width of the distributions between (C) and (D). This is a result of the SMCC diffusion, upon during the transition 2 → 0 (see attached snapshot). Standard errors are plotted for all averaged quantities and in most cases are smaller than the data points.

We also measured the mean captured segment length in the ATP-bound state of the SMC complex (orange points in Fig. 3B). The data show a similar trend to the translocation steps, but are shifted to larger values by an approximately-constant distance. This indicates that DNA slippage occurs in the ADP-bound state, when the SMCC is not bound to DNA at all (transition 2 → 0 in Fig. 2A), by an amount essentially insensitive to DNA tension. The size of the captured segment is comparable to the persistence length of DNA (50 nm) and can be well described by a simple free-energy-minimization model (Supplementary Data, Fig. S2).

Fig. 3C shows the distribution of capture-segment lengths in the ATP-bound state for a few values of DNA tension (all distributions are normalized so that their integrals are 1, hence the units of step size distribution of nm^−1^). We note the appearance of two distinct peaks at intermediate and high forces (*i.e.*, above about 0.2 pN). These peaks are indicative of two distinct substates, the relative population of which is tension dependent. The main distinction between the two substates is the attachment of a DNA segment to the top binding site (see snapshots), which is directly controlled by entropic fluctuations. The attached state is suppressed by high DNA tension, which favors state 1a over 1b. Movies S1 and S2 in Supplementary Data show a typical translocation cycle at low and high DNA tension, respectively, and highlight this distinction.

For comparison, the respective distributions of the translocation step size are plotted in Fig. 3D, and are substantially-wider than their ATP-bound captured segment counterparts, with a tension-independent spread. This suggests the existence of appreciable random diffusion of the SMCC during the cycle, and in particular during the transition from the ADP-bound state back to the apo state (snapshot in Fig. 3D), when the SMCC is not directly attached to DNA. As a result, the SMCC can perform some backward steps (negative tails in Fig. 3D), but it executes a net directed motion after averaging over many whole cycles. The translocation direction is solely determined by the orientation of the SMCC upon its topological loading, since the DNA track is left-right symmetric. The left-right asymmetry, necessary for processive motor activity, arises from the folding of the kleisin protein. Processivity is maintained by the topological linkage between the SMCC and the DNA (Fig. 2A).

### Loop extrusion

Next, we investigated the loop extrusion process by the SMCC model, based on the safety-belt mechanism (Fig. 2B). We distinguish between two cases, depending on the origin of tension in DNA. In the first c ase, t he t ension is externally fixed, b y a pplying a s tretching f orce t o t he ends of the molecule. This can be realized with single-molecule micromanipulation techniques, such as optical or magnetic tweezers. In the second case, it is the end points of DNA that are externally fixed, w hich h as b een experimentally demonstrated *e.g.*, using DNAs attached at both ends to a surface (35, 36, 37). As loop extrusion progresses, the nonextruded DNA is gradually stretched, leading to a corresponding increase in its tension.

### Fixed tension

Figure 4A shows representative time series of extruded loop size during simulations at constant tension, starting from an initial condition of a loop of 400 bp (about 136 nm) captured by the SMCC. The time series are extremely stochastic, and indicate that an initially-loaded loop grows for small tensions, and shrinks for larger ones. Averaging over many such runs leads to a smooth dependence of the mean loop extrusion step size, as a function of applied tension (Fig. 4B), with extruded steps initially becoming smaller with increasing DNA tension, as for translocation (Fig. 3B).

**Figure 4.**
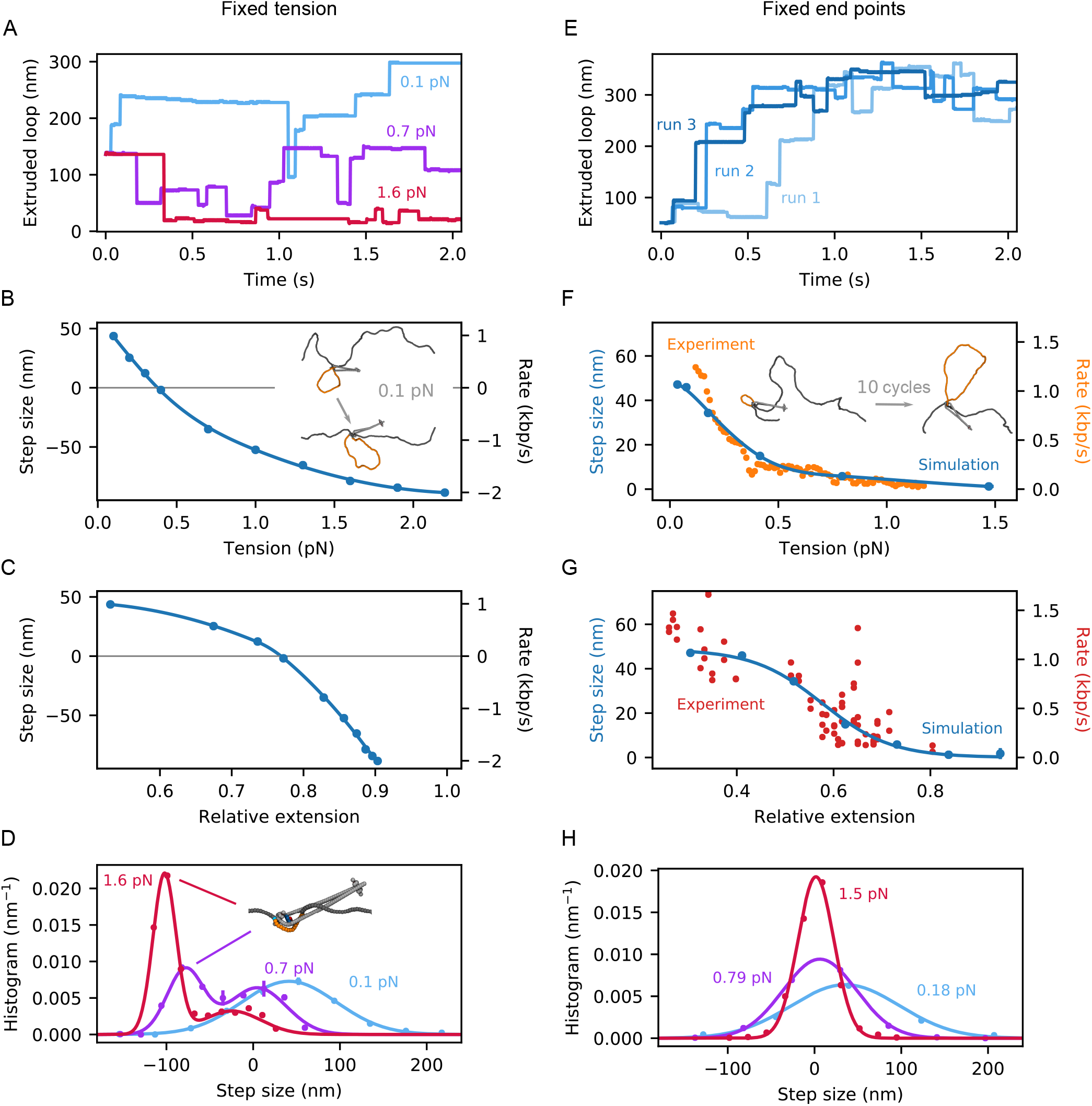
Loop extrusion of DNA by an SMCC under fixed DNA tension (A-D) and end-point distance (E-H). (A) Representative time traces of extruded DNA length by SMC for three selected values of tension. (B) Loop extrusion step size as a function of DNA applied tension. The solid line is a spline interpolation of the simulation data (points). The inset shows the before- and-after configurations for a typical cycle at 0.1 pN. (C) Same data plotted against the corresponding mean relative extension. The solid line is again a spline interpolation of the simulation data (points). (D) Distribution of loop extrusion steps for some selected tensions. The solid lines are single- and double-Gaussian fits to the data (points). The attached snapshot is a representative configuration associated with the second peak at high tensions. (E) Representative time traces of extruded DNA length by SMC for three independent simulation runs. (F) Loop extrusion step size as a function of the mean tension. The latter is computed from the mean net force exerted on the DNA ends. The solid line is a spline interpolation of the simulation data (blue points). For comparison, the experimental data of Ref. (36) are also shown with (orange points). The latter were originally obtained as a loop extrusion rate (right axis), so we transformed them into mean step size (left axis) by multiplying them with a factor of 44.2 s nm/kbp, implying a cycle duration of 0.13 s. The inset shows some typical before- and-after configurations after the lapse of 10 cycles. (G) Same data plotted against the corresponding mean relative extension. The solid line is a fit of a logistic function. For comparison, the experimental data of Ref. (36) are also shown as red points. (H) Distribution of loop extrusion steps for some selected tensions. The solid lines are single-Gaussian fits to the data (points). Standard errors are plotted for all averaged quantities and in most cases are smaller than the data points.

However, unlike translocation, for extrusion, sufficiently large tensions lead to the motor performing backward steps (above 0.4 pN), leading to a net shrinkage of the extruded loop. This takes place during the ADP-bound phase of the cycle (state 2 in Fig. 2B), during which the loop can exchange length with the rest of the DNA. Notably, the amount of slippage per cycle is dependent on the loop size at the beginning of the cycle. A comparison among different initial loop sizes is shown in Fig. S3B of Supplementary Data; for tensions below stalling and slippage, the average step size converges with increasing initial loop size.

Fig. 4C shows the same data, as a function of the mean relative extension. The latter is computed using the known equilibrium relation between force and extension (72). The step size distribution for some selected values of the tension is shown in Fig. 4D, and is bimodal at high tensions. Analysis of the emergent second peaks reveals the presence of “imperfect cycles”: The nontethered part of the DNA does not return to the bottom binding site at the end of the cycle, but instead remains in the upper compartment. This further exposes the loop to the external tension, leading to substantial loop shrinkage, *i.e.*, strongly-negative steps.

### Fixed end-to-end extension

Next, we performed molecular dynamics simulation of loop extrusion at fixed end-to-end DNA extension, as studied experimentally (36). We started these simulations at a relative extension (ratio of end-to-end distance divided by DNA contour length outside the SMCC) of 25%, corresponding to a DNA tension of roughly 0.04 pN. The SMCC performed a total of 10 loop-extrusion cycles per simulation run (see Movie S3 of Supplementary Data). Fig. 4E shows a few representative extruded loop sizes versus time and one sees much less noisy behavior than in the fixed tension case (compare Fig. 4A), with saturation of the extruded loop size at roughly 300 nm (900 bp) size as DNA tension rises to the point where it limits capture of loops by the enzyme.

Fig. 4F shows the mean loop extrusion step size as a function of the average tension (blue points). This is obtained by a direct computation of the DNA tension after each cycle, and subsequent binning of the data. Note that, contrary to the fixed-tension case (Fig. 4B), fixing the end points puts a hard limit on the DNA length that can be extruded, and loop extrusion halts at high tension with a large extruded loop, without reversal. The value of the mean step size at low tension (47 nm per cycle) is in quantitative agreement with that for the fixed-tension case (Fig. 4B).

Ganji et al. have experimentally measured the rate at which condensin extrudes loops in DNA with fixed end points, using a single-molecule assay (36). These experiments were used to set the timescales for all of our simulations, using the fact that the ATP hydrolysis step (1b→2 in Fig. 2) is rate limiting in our simulations, with the other steps being in pre-equilibrium. This allowed us to rescale our cycle time so as to put our simulation and the experimental data on the same time scale. We found that choosing a cycle duration of 0.13 s resulted in an excellent correspondence beween our simulations and experimental data (orange points in Fig. 4F). Figure 4G shows data from the same simulations and experiments, plotted versus relative DNA extension (end-to-end distance over nonextruded DNA length). Again we find good agreement between simulation and experiment, together with the gradual slowing down of the motor at large DNA extension (high tension). This single time rescaling is used to set all the timescales for our MD results in Figs. 3 and 4.

Fig. 4H shows the distribution of the mean loop extrusion step size for some selected mean tensions. In all cases, the data can be well fitted with a single Gaussian peak. Comparison between Figs. 4D and 4H further highlight the equivalence of the two situations (*i.e.*, fixed tension and end-to-end extension) for low DNA tension/extension. Recent observations of step size distributions for yeast condensin (53) are in good agreement with our results.

### Varying the model parameters

In order to investigate how the behavior of the motor depends on the model details, we performed additional translocation, fixed-tension and fixed-end-point loop extrusion simulations for different model parameters (Fig. 5, panels A, B and C, respectively). More specifically, we deactivated the top binding site throughout the whole cycle, and found that the motor could still operate, suggesting that the existence of the top binding site is not a necessary element in the model (blue points). Interestingly, the translocation of the motor was found to be more efficient at high tension. This is likely due to the 2 → 0 transition being faster in that case, since DNA needs to travel a shorter distance until it reaches the bottom binding site. Similarly, the SMC complex was found to be insensitive to a deactivation of the middle binding site (red points).

**Figure 5.**
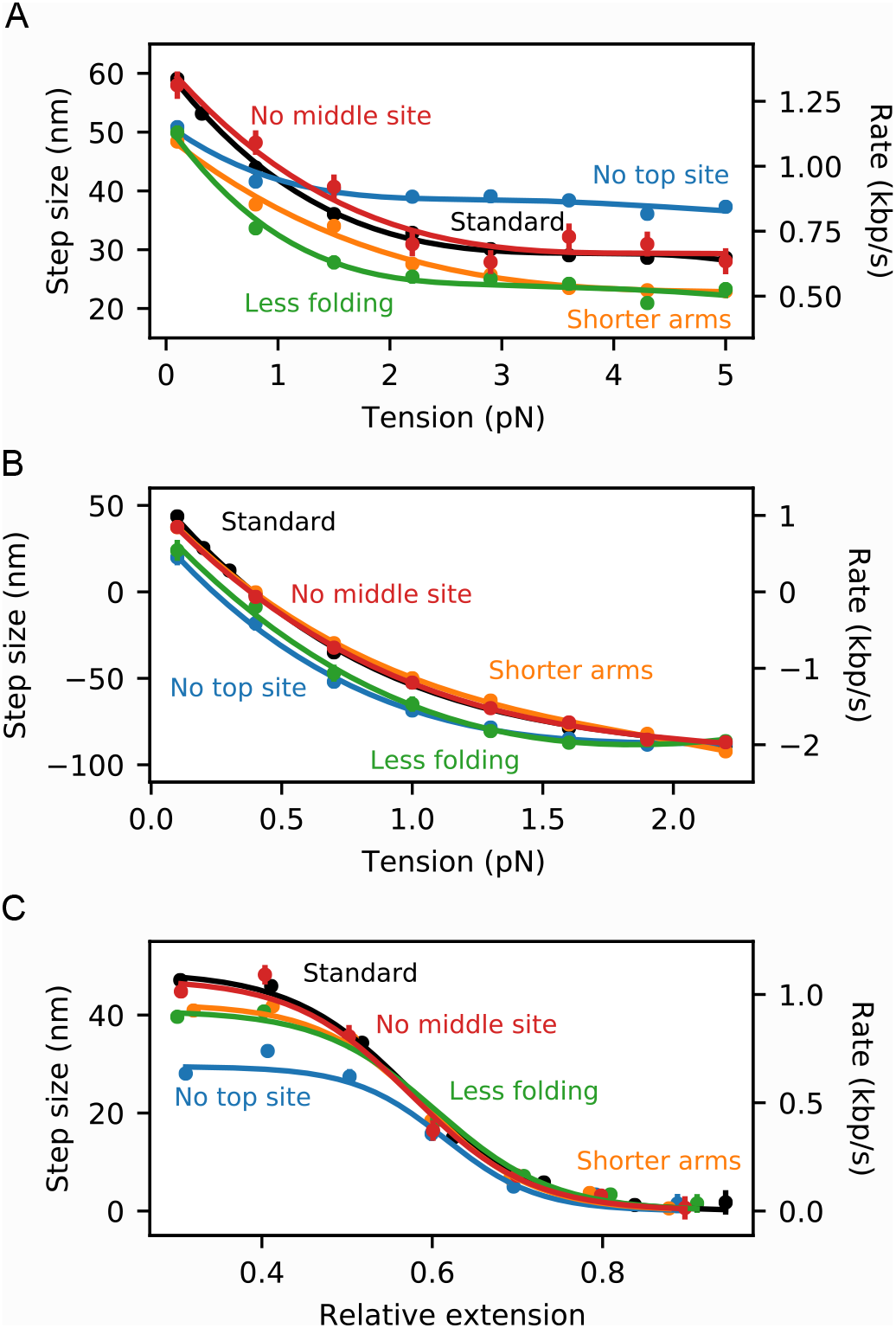
Comparison among different sets of model parameters for (A) translocation, (B) loop extrusion under fixed tension and (C) loop extrusion under fixed end-point distance. The black points correspond to the parameters used throughout this work (standard), the blue points to a deactivation of the top binding site (no top site), the red points to a deactivation of the middle binding site (no middle site), the orange points to a shortening of the coiled-coil arms from 50 nm to 40 nm (shorter arms) and the green points to a reduction of the folding angle of the lower compartment in the ATP-bound state from 160 degrees to 130 degrees (less folding). Standard errors are plotted for all averaged quantities and in most cases are smaller than the data points.

Additionally, we reduced the length of the coiled-coil arms from 50 nm down to 40 nm, and observed little to no difference (Fig. 5, orange points). We also reduced the folding angle of the lower compartment in the ATP-bound state from 160 to 130 degrees, which slightly slowed down the motor but did not keep it from translocating (green points). Finally, making the elbows in the SMC arms completely flexible (zero bending stiffness) did not keep translocation from occuring, although it proceeded via somewhat smaller steps at low force (Fig. S3A, Supplementary Data). This emphasizes that conformational and mechanical details of the upper compartment are less important than the breaking of symmetry and topological separation of upper and lower compartments that occurs upon ATP binding. We conclude that the MD model motor function is qualitatively insensitive to a variety of modifications, and that it is likely applicable to a range of SMCCs. The quantitative predictions of the specific modifications we have examined amount to predictions for realizable SMCC mutation experiments.

## DISCUSSION

The results presented above comprise a detailed analysis of the segment-capture model for SMC translocation and loop extrusion (Fig. 2) (19, 41). The numerical nature of our model circumvents analytical limitations (41), and allows us to make a number of predictions for future experiments.

### Translocation

Our model predicts the DNA-tension dependence of translocation by SMCCs (Fig. 3B), with the translocation rate dropping as DNA tension is increased through about 1 pN. Prior analytical modeling (41) did not fully explore the effect of a possible power stroke, in the form of kleisin folding. In our MD simulations, this allows translocation to proceed even when the DNA is tightly stretched, via an “inchworm”-like mechanism (smaller peak in captured segment size distribution, Fig. 3C). The drop to a plateau translocation rate with increasing force (Fig. 3B) is the signature of translocation occurring through two distinct mechanisms (two peaks in Fig. 3C), namely DNA bending and segment capture at lower forces *vs.* “inchworming” (possibly involving SMC bending) at high forces. We note that substrate-tension-feedback control of SMC function has been seen in other models (73), an effect similar to our observation of a change in translocation mechanism as DNA tension is varied.

To date, no translocation experiments at controlled tension have been carried out. Observation of translocation by yeast condensin observed a velocity of 60 bp/s for DNA tension estimated to be 0.3 pN (35). This is substantially slower than the respective estimate of about 1.2 kbp/s from our MD model (Fig. 3B), the cycling rate of which was set using loop-extrusion experiments (36). This deviation may, thus, reflect a large variability in the motor efficiency, depending on the precise experimental conditions.

In addition to the tension-velocity behavior, the MD model makes clear predictions for the size distribution of the DNA segment capture (Fig. 3C) (which, for the inchworm mode, corresponds to essentially the length of the SMCC), as well as for the translocation step (Fig. 3D). The model shows that the relatively-narrow captured DNA segment size is broadened by diffusion into a more smeared-out DNA step size distribution. The smearing is a result of the entropic transport of DNA from the top binding site back to the bottom one following ATP hydrolysis (Fig. 2 2 → 0), during which diffusion of the SMCC along the DNA can freely occur. Cohesin has been observed to diffuse on DNA (74) in experiments lacking the motor-processivity protein NIPBL (37).

The model’s key transition, which breaks left-right symmetry and provides a power stroke, is head engagement and folding of the lower compartment (Fig. 2 0 → 1a), guiding the DNA to the middle binding site on top of the Walker ATPase heads (Fig. 2 1a). This leads to formation of a DNA segment in the upper compartment (Fig. 2 1a ↔ 1b). A remarkable result is that the top binding site is dispensable, in that translocation and loop extrusion persist without it (Fig. 5A-C, blue curves). In accord with this, a recent genetic experiment on bsSMC mutated away putative DNA-binding residues in the upper “hinge” domain, with little effect on translocation and loop extrusion *in vivo* (25). We also have observed that the middle binding site on the ATPase bridge is also not required for translocation and loop extrusion (Fig. 5A-C, red curves).

Our observation of a lack of necessity for the upper-compartment DNA-binding site may help to explain the variability in the hinge-proximal DNA-binding across SMCCs. While the presence of a positively-charged “channel” (with, so far, poorly-understood function) along with a positively-charged surface on the hinge are conserved (57), the precise location of basic residues on the hinge surface varies appreciably across SMCCs (55, 56, 59, 60). This variability may reflect our result that the top binding site is not crucial, and is involved in fine-tuning of SMCC function. Our model enzyme cycle does require DNA release from the top site following arm refolding, and we can expect the top site to be relatively weak, so as to have its interaction with DNA disrupted by the return to the apo state.

Current experiments and Fig. 3 do not consider the effect of a load force, examining DNA translocation only as a function of DNA tension. It would be informative to additionally apply a direct load to the enzyme, and to examine translocation velocity as a function of both DNA tension and enzyme load force. This could be realized with a combination of three applied forces using, *e.g.*, triple force-controlled optical tweezers, with forces *f*_tension_ and *f*_tension_ − *f*_load_ at the two ends of the DNA, and *f*load applied to the SMCC (41). A recent study observed translocation for cohesin at a velocity of 0.4 kbp/s against a buffer flow (37). The latter may introduce both DNA tension (by stretching) and a load force on the SMCC, although in an imperfectly quantitatively-controlled fashion.

### Loop extrusion: fixed end points *vs.* fixed tension

The MD model quantitatively describes compaction rates observed in experiments on loop extrusion at fixed DNA end-point distance (36) (Fig. 4F,G), and additionally predicts step sizes and their distributions (Fig. 4H). The tension in the DNA builds up as a loop is extruded, and there comes a point at which the enzyme stalls. Due to this externally-induced tension-extrusion coupling, the loop can never shrink, and thus the MD rate *vs.* tension is always positive, asymptotically approaching zero for large forces, as observed experimentally (Fig. 4F,G).

The positivity of compaction rate with fixed end points is in stark contrast to the situation for fixed DNA tension, where there is a well-defined stall tension, beyond which an initially-extruded loop will start shrinking (Fig. 4B,C). Indeed, our MD model is eventually forced to run in reverse, taking negative steps of well-defined s ize ( Fig. 4 D). E vidence f or reversal of SMCC loop extrusion/translocation by force exists, in the form of Hi-C data from *B. subtilis* consistent with bsSMC-RNAP collisions forcing bsSMC backwards along DNA (75). Experiments on loop extrusion *vs.* controlled DNA tension would provide further insight into how SMCCs on DNA can be pushed around by other enzymes. *In vivo* one can imagine loop extrusion being opposed by both fixed endpoint and fixed tension restraints, the former being relevant to a chromosomal domain, which is defined by binding to solid cellular structures at two distant points, and the latter being relevant to molecular motors, such as polymerases, which might act to generate tension in a DNA segment.

### Head engagement power stroke

In the MD model, ATP binding and head engagement are associated with a conformational change of the SMCC, which facilitates segment capture from one side of the enzyme (Fig. 2 0 → 1). Such a symmetry-breaking event is necessary for the translocation to be directional, otherwise DNA segments would be captured with equal efficiency from both directions, and the enzyme would move randomly left and right along its unpolarized DNA track. This conformational change might be directly observable in an experiment that monitors the enzyme itself, *e.g.*, by monitoring the distance between ATPase heads directly, or the effect of head engagement on overall conformation of the enzyme.

In our SMCC model, head engagement and kleisin folding move the lower edge of the enzyme by a distance of roughly 10 nm (vertical distance moved by lower edge between states 0 and 1 in Fig. 2 0 → 1). In our model, this transition can actually be observed in terms of a small contraction in the flanking DNA (Fig. S4, Supplementary Data), although to actually observe this it is likely that a quite short DNA will have to be used (our MD simulations use a total of 1.5 kb).

By applying sufficient force against this conformational change, one might be able to keep it from occurring, providing measurement of the enzyme power stroke. The threshold to overpower the conformational change would likely be a force in the vicinity of 10 pN, *i.e.*, the force scale associated with breaking noncovalent biomolecule-biomolecule interactions, or the stalling of molecular motors. The origin of this force scale is in the range of forces required to change conformation of or to unfold proteins by force; in this case, one is acting against the force driven by the binding of ATP to link together the two Walker ATPase subunits. This force can be estimated by dividing the free energy associated with ATP hydrolysis (≈20 *k*_B_T) per ATP) by the conformational change (≈3 nm), which leads to ≈25 pN. Our results show that this force is associated with the stalling of translocation but is only indirectly related to that for loop extrusion (Fig. 3B *vs.* Fig. 4F). Recent observations of large ATP-hydrolysis-independent contractions of SMCC-DNA complexes (53) are likely looking at the DNA segment capture process rather than protein conformational change.

### SMCC conformation and flexibility

A feature of the model that we have explored is the effect of SMC coiled-coil “arm” flexibility on SMCC translocation and loop extrusion. We have found that making SMC arm joints completely flexible does not abrogate translocation (Fig. S3A, Supplementary Data), and in fact eliminates the need for DNA bending, thus likely facilitating translocation at high DNA tensions. This may be important, given that there are observations of rather extreme SMC arm flexibility for condensin (47, 76), although recent cryo-EM images of precisely the same type of condensin (20) suggest conformational properties similar to what we have assumed here. However, essentially the same motor function should result from variations on the protein-ring closure mechanism presumed here, for example an ATP-dependent “collapse” of the protein ring (53).

The main features needed for SMCCs to translocate in the manner described by our model are the folding/breaking of symmetry in the lower compartment, and the formation of separated upper and lower compartments, both a result of ATP binding. The key elements of our model are largely topological: passage of DNA through the protein ring, and the ATP-dependent division of the ring into upper and lower compartments. Models of SMC function which do not incorporate the topological nature of SMCC-DNA interaction will have weak directional processivity due to the left-right symmetry of duplex DNA. Given this, changes to SMCs which reduce the area of opening of the coiled-coil upper compartment will likely tend to impairment of translocation and loop extrusion.

We have found that DNA binding interactions, apart from some mechanism to hold on to DNA in the lower compartment, are largely dispensable, with elimination of the upper and middle sites not significantly changing translocation and loop extrusion (Fig. 5). Our model may overestimate this robustness due to the strong symmetry-breaking folding of the kleisin, which allows MD simulation of loop capture on a computationally-manageable timescale, but which may also lead to an underestimation of the importance of the loop-capturing DNA binding sites. Modification of the strength of the DNA binding sites is also a reasonable way to model changes in univalent salt concentration, with weaker interactions corresponding to higher salt concentration. However, precise quantitative calibration of this in our coarse-grained description is problematic.

### Loading topology and symmetry of loop extrusion

As discussed earlier, translocation is the fundamental function of SMCCs that underlies all modes of loop extrusion. Depending on the SMCC, it appears that different modes do occur, with distinct loading mechanisms and symmetry of loop extrusion (Fig. 6). For example, it has been suggested that yeast condensin translocation and loop extrusion require passage of the DNA through a transient opening of the SMCC protein ring (33). For this SMCC, the apparent anchoring of the DNA to the outside of the ring (safety belt) requires topological loading, and leads to asymmetric loop extrusion (Fig. 6A), formally the mode used in our MD model.

**Figure 6.**
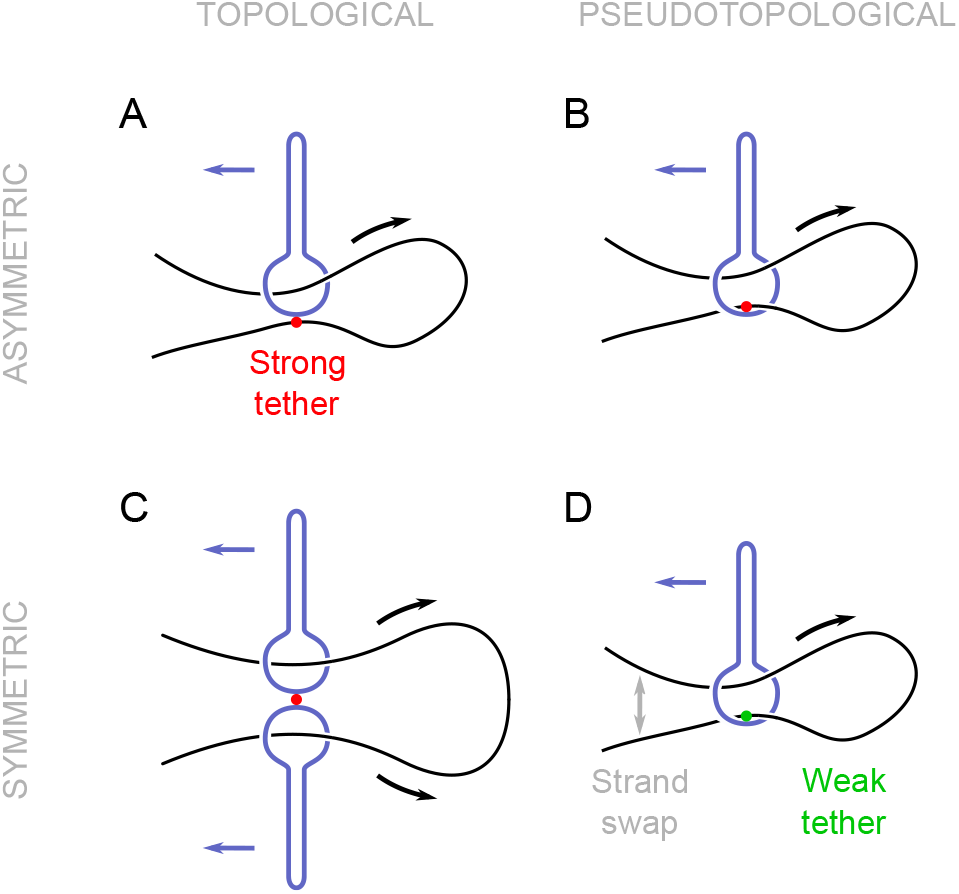
Different loop extrusion mechanisms, categorized according to loading mechanism (vertical) and symmetry of loop extrusion (horizontal). Permanently attaching DNA to the exterior of the SMCC (safety belt, marked with red) leads to asymmetric loop extrusion. This requires DNA threading through the lower compartment, and is the mechanism used in this work. (B) Moving the tethering point to the interior of the SMCC does not require opening of the protein ring, but rather an insertion of a DNA hairpin. (C) Two permanently-joined SMCCs translocating in opposite directions can symmetrically extrude loops. (D) If the tether in the interior of the SMCC is weak, such that DNA detachment and strand swap can take place, the same pseudotopological model as in (B) can now symmetrically extrude loops.

In principle, our MD model may also be applied to the case where the DNA is bound *inside* the SMC-kleisin ring (Fig. 6B). While this is geometrically identical to the external safety-belt scheme (the relocation of the tethered DNA strand from the outside to the inside of the kleisin is inconsequential to the cycle of Fig. 2B), there is a key topological difference: no opening of the protein ring is required to initiate the loop-extrusion process, since the DNA can be bent into a hairpin and then inserted into the SMC-kleisin ring (pseudotopological loading) (41). As long as DNA remains tethered in the interior of the SMC-kleisin ring, this mechanism generates one-sided, asymmetric loop extrusion, similar to the external safety belt. Recent DNA mapping studies suggest that condensin may bind DNA in this manner (66).

A third possibility is a simple variation of the above, where the tethering of DNA is weak enough to allow transient detachment and reattachment (Fig. 6D). Since two DNA segments are in close proximity in the interior of the SMCC protein loop, a strand swap can take place, possibly via facilitated dissociation (likely a strong effect under the strong confinement of two DNA strands in the SMCC lower compartment) (77). If this occurs relatively slowly compared to the cycling time, the result will be progressive extrusion on both sides of the loop *i.e.*, symmetric pseudotopological loop extrusion, as indeed has been observed for cohesin (37). Each strand swap event translates to a change in which side of the loop DNA segments are captured from, which might be detectable given sufficient temporal resolution.

Finally, a fourth possible “dimerized translocator” can result from the coupling of two SMCCs together (Fig. 6C), each translocating to an opposite direction, so as to accomplish symmetric loop extrusion at double the velocity of a single translocator (41). This scenario requires topological loading of DNA segments into both of the dimerized SMC-kleisin rings. Evidence exists for dimerization of the *E. coli* SMCC MukBEF *in vivo* (78, 79, 80), as well as for ATP-dependent compaction activity by oligomerized condensins *in vitro* (34, 38). Examination of all of these scenarios for loop extrusion using MD simulations of the sort we have reported here would be desirable. However, they all present a higher degree of computational difficulty than that of the current paper, with the dimerized translocator being particularly extreme in this regard.

This suggests a rule: single SMCCs that require topological loading *must* asymmetrically extrude loops (at least between successive protein-loop-opening events), while ones that load pseudotopologically *can* perform symmetric loop extrusion (cohesin). A recent study indicates that condensin from metaphase *Xenopus* egg extracts drives more asymmetric loop extrusion than does cohesin from interphase extracts (39), consistent with condensin being topologically loaded, and (interphase) cohesin being pseudotopologically loaded. The precise rules for how loading and loop extrusion occur *in vivo* are likely regulated by factors that mediate SMCC protein loop opening (62, 65).

### Obstacle bypass and formation of z-loops

SMCCs have recently been observed to be able to “bypass” an obstacle during loop extrusion, in the sense that loop extrusion has been observed to continue following the encounter of obstacle and SMCC (81). This appears inconsistent with the long-established topological linkage between DNA and tripartite SMCC protein rings (22, 23), and with segment-capture-models (19, 41) which incorporate this feature to obtain processivity, translocation and loop extrusion (19, 41). However, a resolution of this apparent conflict has been proposed for the segment-capture model (66) (note that Fig. 6 of Ref. (66) is topologically equivalent to the variant of loop extrusion for the segment-capture model shown in Fig. 7B in Ref. (41) and Fig. 6B of this paper, all with an initial loop of DNA passing twice through the lower compartment, a subsequent ATP-binding-driven “power stroke” guiding upstream DNA segment capture in the upper compartment, and then ATP hydrolysis and Walker ATPase separation which causes enlargement of the initial DNA loop by merging it with the captured DNA segment (82)).

In the new experiments (81), a relatively large colloidal particle was tethered to a specific point along the DNA, leading to a collision between an SMCC and the obstacle. Fig. S6A-B illustrates this for translocation: when all the DNA “downstream” of the obstacle (Fig. S6A, purple) is pulled through the SMCC, a point is reached where the obstacle blocks further translocation (Fig. S6B). The key insight of Ref. (66) is that in this obstacle-blocked state, DNA on the *upstream* side of the obstacle (Fig. S6, green DNA) can still be captured by the SMCC leading to extrusion of a loop with the obstacle effectively acting as the “loop anchor” (Fig. S6C-F). In the experiments of Ref. (81), the use of a flexible PEG linker between the DNA and the large bead-obstacle may facilitate this process, but similar “obstacle bypass” can likely occur past some objects that are directly bound to the DNA, so long as the downstream DNA is able to be captured by the middle (bridge) binding site. A consequence of this that is biologically relevant is that obstacles that are able to block SMCC translocation may act as loop-extrusion initiation sites, causing compaction of downstream DNA into an extruded loop.

Fig. S6G-H show this process including anchoring of the upstream (purple) DNA to form a loop (loop 1) which cannot grow beyond the obstacle attachment point. Capture of the downstream DNA (green) leads to formation of a second loop (loop 2). In this state, if one applies a force to the obstacle (e.g., via fluid flow in the case where the obstacle is a relatively large colloidal particle) the topology is such that a third loop can be “pulled out” of the SMCC (Fig. S6H, loop 3).

The same general ideas can be applied to the case where the obstacle is a loop-extruding SMCC along the same DNA as a second loop-extruding SMCC (Fig. S7A). As the “outer” SMCC (right, condensin b) approaches the “inner” one (left, a), DNA downstream (red) of the inner SMC can be captured in the outer one (Fig. S7B), leading to extrusion of a new loop of downstream DNA (Fig. S7C). When rearranged without topology change, this is easily deformed into a “z-loop” structure as observed in recent experiments with colliding SMCCs (Fig. S7D)(83). These structures can be undone using force (e.g., fluid fl ow), by pu lling th e ne wly ex truded loop back out of the z-loop (Fig. S7E). This general scheme is also consistent with Hi-C maps indicating traversal of one SMCC by another *in vivo* which have been interpreted in terms of z-loop formation (84).

## ACKNOWLEDGEMENTS

The authors acknowledge helpful discussions with Dr. Christian Haering.

## FUNDING

This work was supported by the Research Funds Flanders (FWO Vlaanderen) [grant number VITO-FWO 11.59.71.7N, S.K.N.] and by the National Institutes of Health [grant numbers R01GM105847, U54CA193419 and UM1-HG011536, J.F.M].

## Conflict of interest s tatement

The authors declare no conflicts of interest.

## Data availability

The LAMMPS parameters that define the MD model and the scripts used to generate simulation data are available at https://github.com/sknomidis/SMC_LAMMPS.

All numeric data are available on request to the corresponding author.

## Supplementary Data

### Supporting Information Text

#### DNA binding sites

**Table S1.** DNA binding sites assumed in model along with corresponding protein regions and supporting experimental data. Left column lists first the model nomenclature (top, middle, bottom, anchor/safety belt) and the related structural features. Middle column lists protein subunits, amino acid residues involved in DNA binding, and relevant references, for the *B. subtilis* SMC. Right column lists the protein subunits, residue numbers, and references for yeast condensin.

| DNA binding site |  | Bsu Smc-ScpAB |  |  | Yeast condensin |  |  |
| --- | --- | --- | --- | --- | --- | --- | --- |
|  |  | Subunit | Residues | Reference | Subunit | Residues | Reference |
| top | hinge | Smc | K666, K667, K668 | (1) (2) | Smc2<br>Smc4 | positively charged<br>surface patch observed | (3) (4) (5) (6) |
| middle | ATP-heads | Smc | R54, K60, R120<br>K122, R153 | (2) | Smc2<br>Smc4 | site likely based on wide<br>conservation (Rad50 and *Bsu* Smc) | (2) (7) |
| bottom | kleisin-kite/hawk | ScpAB | Steric binding (putative) |  | Ycs4 | site likely based on<br>similarity to cohesin Scc3 | (8) |
| anchor | safety-belt |  | Unknown |  | Ycg1<br>Brn1 | K70, K71, K849<br>K409, R411, K456 | (9)<br>(9) |

#### Coarse-grained SMCC model

**Table S2.**
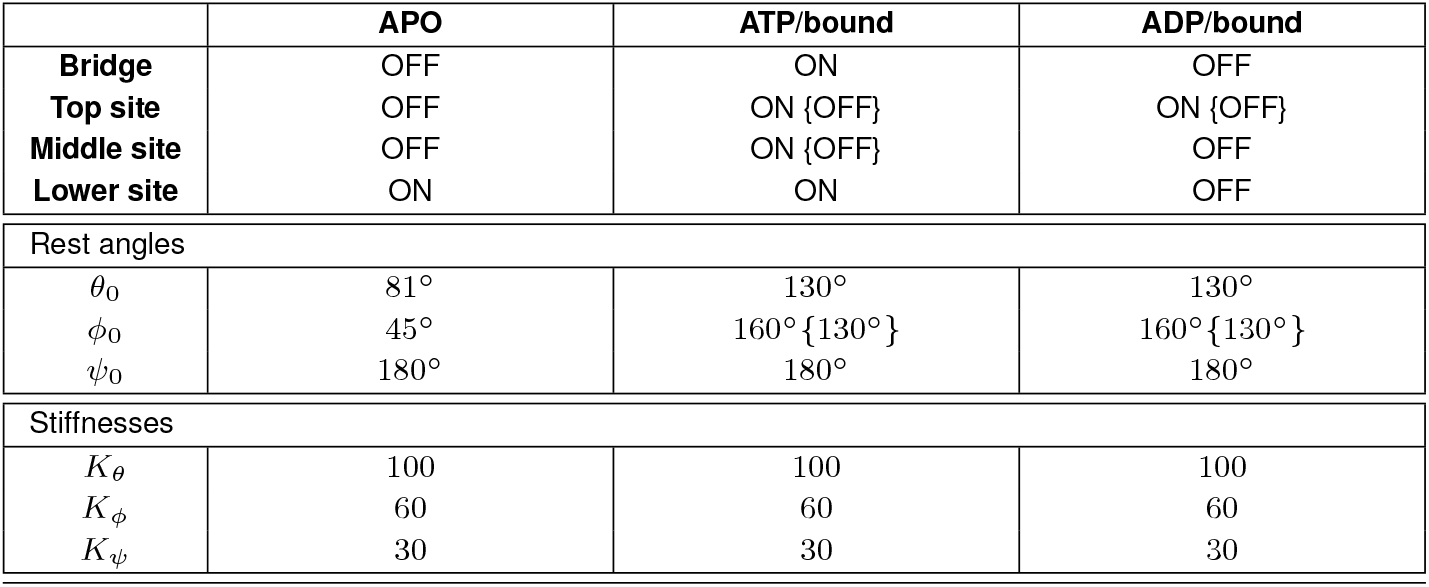
Summary of the SMCC simulation parameters used for modeling the three different states. The angles are defined as shown in Fig. S1 and are expressed in degrees, while stiffnesses in units of *k*_B_*T*/rad^2^. The curly brackets indicate the values used to produce some of the variations of Fig. 5 of the main text.

|  | APO | ATP/bound | ADP/bound |
| --- | --- | --- | --- |
| Bridge | OFF | ON | OFF |
| Top site | OFF | ON {OFF} | ON {OFF} |
| Middle site | OFF | ON {OFF} | OFF |
| Lower site | ON | ON | OFF |
| Rest angles |  |  |  |
| $\theta_0$ | 81° | 130° | 130° |
| $\phi_0$ | 45° | 160° {130°} | 160° {130°} |
| $\psi_0$ | 180° | 180° | 180° |
| Stiffnesses |  |  |  |
| $K_\theta$ | 100 | 100 | 100 |
| $K_\phi$ | 60 | 60 | 60 |
| $K_\psi$ | 30 | 30 | 30 |

#### Captured segment length

The translocation and loop-extrusion function of the SMCC is based upon a segment-capture process, which takes place during its ATP-bound state (Fig. 1C,D of main text). Here, we present an estimate of the segment length *vs.* the DNA tension, based on a free energy minimization. We assume that the SMCC stays sufficiently long at that state, so that we can treat the system as being at equilibrium. As the captured segment length (Fig. 2A of main text) is comparable to the persistence length of DNA (50 nm), one may neglect the former’s conformational fluctuations, and focus on its minimal-energy shape (below we refer to the bent DNA segment captured during a translocation cycle as a DNA “loop”; we note that this is distinct from the larger *extruded* loop). The total free energy of the system is then given by

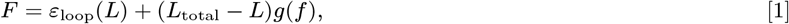

*L*_total_ is the total DNA length, *ε*_loop_(*L*) the minimal energy for a loop of length *L* (blue line in Fig. S2A) and *g*(*f* ) the free energy per unit length of a linear (*i.e.*, unlooped) DNA under a tension *f* (black line in Fig. S2A). To obtain the loop length for the given DNA tension, one needs to minimize Eq. (1) with respect to *L*. Note that, we have omitted length-independent terms in Eq. (1), as they do not affect our calculation.

Both *g*(*f* ) and *ε*_loop_ can be estimated based on the wormlike chain model, which treats DNA as a semiflexible polymer. The free energy density of a stretched DNA can be obtained by integrating its relative DNA extension with respect to the tension. In the high-force limit, this is simply (10)

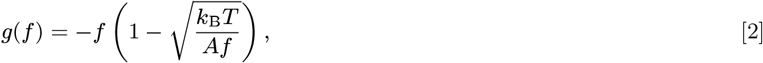

where *A* = 50 nm is the persistence length of DNA and *k*_B_*T* ≈ 4.1 pN nm is the thermal energy at room temperature. Since, here, we are also interested in intermediate forces (down to 0.1 pN), we will numerically estimate *g*(*f* ) using an interpolation formula for DNA extension (11), see Fig. S2C.

The loop can be parametrized with an angle *θ*, denoting the orientation of the tangent with respect to the *x*-axis (Fig. S2A), while its energy is given by

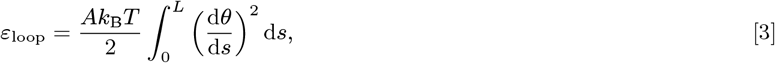

where *s* is the arclength coordinate. We wish to minimize this energy with respect to *θ*(*s*), under the constraints *θ*(0) = *π*/2 and *θ*(*L*) = 3*π*/2, *i.e.*, by keeping the tangents antiparallel at the loop ends points, which resembles the action of the SMCC on DNA in the ATP-bound state (state 1b in Fig. 1C,D of main text). A simple variational solution can be obtained through a circle-line approximation (12, 13), as illustrated in Fig. S2B. Loops of length *L < πd*/2 (with *d* the end-point distance) are approximated with two quadrants connected with a straight line, whereas longer loops by a semicircle extended with straight lines. The loop energy of such a circle-line model is

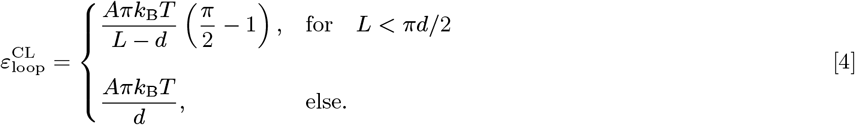

The loop shape can also be obtained from a Fourier-series expansion of the curvature, d*θ*/d*s*, as recently shown in Ref. 14 for closed and open loops with free boundary conditions. It turns out that, under fixed boundary conditions, *θ*(0) and *θ*(*L*), the Fourier series can be rewritten into the simple form (unpublished result)

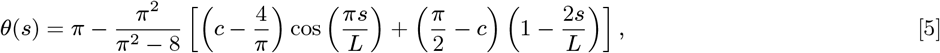

where the parameter *c* fixes the relative end-point distance, *d/L*, as follows

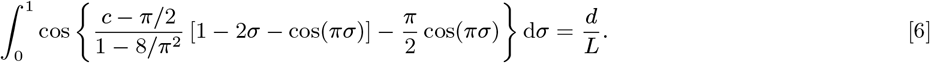

Note that, *c* = 4*/π* describes a semicircle, while *c* = *π*/2 a loop with a single Fourier component. Finally, substitution of Eq. (5) into Eq. (3) yields the energy

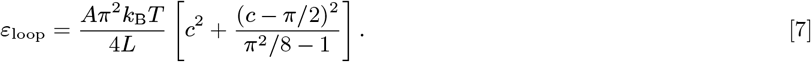

Figure S2D shows a comparison between Eq. (4) and Eq. (7) (dashed and solid lines, respectively), for varying loop size. Obviously, the exact calculation yields consistently-lower energies than the circle-line approximation, with the exception of the semicircle case, *L* = *πd*/2, where the two coincide. Moreover, smaller loops have a larger energy, due to their increased curvature. Note that, the circle-line energy is constant for *L > πd*/2, as any additional loop length is absorbed into the energetically-inconsequential straight segments.

One can now minimize Eq. (1) with respect *L*, using either Eq. (4) or Eq. (7), so as to obtain the minimal-energy loop size for fixed DNA tension, shown in Fig. S2E with dashed and solid green lines, respectively. For comparison, the loop-size MD data from Fig. 2A of main text are shown (blue points). At high DNA tension, the theory deviates from the simulation data, as partial segment capture starts taking place (state 1a in Fig. 3C of main text), which is not accounted for by this theory. Excluding these events from the calculation of the mean step size (orange points), reveals a good agreement with the theory. Note that, the loop end-point distance is the only parameter of the calculation, and it is fixed at *d* = 35 nm from the SMCC geometry.

#### DNA contraction during the SMC cycle

In the SMCC model, the upward folding of the kleisin moves the lower edge of the enzyme by a distance of approximately 10 nm. This is the vertical distance moved by lower edge when a transition between the states 0 and 1 takes place (see Fig. 2 of the main text). These transitions can actually be observed in simulations as small contractions in the flanking DNA, as illustrated in Fig. S4. When the SMCC is in the state 1, the end-to-end distance of the DNA is shorter than with SMCC in states 0 or 2. The difference is small and decreases at higher applied tension.

#### Testing the overdamped dynamics of DNA

In coarse-grained simulations with implicit solvent, the mass assigned to each coarse-grained bead should also take into account the correlated motion of the surrounding fluid. LAMMPS (15) integrates the equations of motion using a total force of the type *F* = *F_c_* + *F_f_* + *F_r_* with *F_c_* a conservative force, *F_f_* = −*mv*/d_m_ a viscous force and 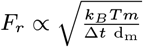 a random force. In the previous expressions *v* is the velocity, dm is the damping coefficient and ∆*t* the integration timestep. The viscous and random forces depend on the mass through the ratio *m*/d_m_, while the mass enters in the equations of motion *ma* = *F* through the inertial term *ma*. Inertia has typically an effect on the short time scale dynamics, while it can be safely neglected at long times. In order to probe inertial effects we performed some test simulations on a our coarse-grained model of DNA by rescaling the mass and damping coefficients by the same factors so to keep the ratio *m*/dm constant. We considered a loop of length *L* = 136 nm (400 bp) with fixed endpoint and probed the dynamics of the central monomer for which we calculated the mean squared displacement ∆*r*^2^(*t*). Figure S5 shows plots of this quanitites for various values of bead mass of DNA, with *m* being the value reported in the main text. The excellent overlap of the data indicate that inertial effects are negligible beyond the timescale of 1 ns.

#### Obstacle bypass by SMCC

In a recent experimental study (16) a nanoparticle of about 200 nm in size was attached to a specific location on DNA via a PEG-linker. The nanoparticle is much larger in size than a SMCC protein ring ( 50 nm). Loop extrusion was then observed to not necessarily stop when the SMCC pulls DNA in to the point where the loop base and SMCC meet the large obstacle. This observation was interpreted as proving that the SMCC can “bypass” the obstacle and therefore that the DNA cannot be inside the (crosslinked) tripartite protein ring. The observations are done using optical microscopy, with a maximum imaging resolution in the 100 nm range and one observes obstacle and DNA loop (bright dot indicating a random coil) in the same location, to optical resolution. The experiment cannot test whether the obstacle has passed through the protein ring. In the case that our model SMCC encounters such an impassable obstacle, our segment-capture mechanism can continue to operate as illustrated in Fig. S6. In conclusion, we believe that the recent experiments (16) do not invalidate the segment capture model discussed in this paper. It is certainly interesting to investigate quantitatively how the present SMCC model copes with obstacles of various sizes. This is beyond the scope of the present manuscript and will be left for future studies.

#### Condensins crossing one another

Recent experiments on interactions condensin-DNA showed, quite surprisingly, that two condensins can traverse one another creating so-called Z-loops, where three double-stranded DNA helices align in parallel with one condensin at each edge (17). Figure S7 shows a possible explanation of this phenomenon within the segment-capture model.

#### Summary of statistics used to obtain averages

All data was derived from simulations of strings of SMCC ATP cycles, which we refer to as “steps”. Each step consisted of an average of 2.4 *μ*Lsec of simulation time (0.4 in the apo state, 1.6 in the ATP state, and 0.4 in the ADP state). Since the microscopic MD timestep was 0.2 pLsec, 1.2 10^7^ MD steps were used to simulate each SMCC cycle (SMCC step).

**Figure 3:** 7997, 11226, 7838, 7599, 7659, 7826, 7500, 7891, 7821 SMCC steps were used for 0.1, 0.32, 0.8, 1.5, 2.2, 2.9, 3.6, 4.3, 5 pN, respectively.

**Figure 4A-D:** 996, 1337, 1345, 1004, 965, 1069, 1057, 964, 905, 903 steps were used for 0.1, 0.2, 0.3, 0.4, 0.7, 1, 1.3, 1.6, 1.9, 2.2 pN, respectively.

**Figure 4E-H:** A total of 35806 steps were used. These were binned as: 7395, 6767, 5609, 6471, 6785, 2640, 139 steps were used for relative extension 0.3, 0.41, 0.52, 0.62, 0.73, 0.84, 0.94, respectively.

**Figure 5A red:** 910, 989, 1024, 1032, 971, 944, 996, 957 steps were used for 0.1, 0.32, 0.8, 1.5, 2.2, 2.9, 3.6, 4.3, 5 pN, respectively.

**Figure 5A blue:** 4183, 4115, 4124, 4368, 4463, 4160, 4235, 4172 steps were used for 0.1, 0.32, 0.8, 1.5, 2.2, 2.9, 3.6, 4.3, 5 pN, respectively.

**Figure 5A green:** 3116, 3304, 3376, 3487, 3443, 3445, 3370, 3408 steps were used for 0.1, 0.32, 0.8, 1.5, 2.2, 2.9, 3.6, 4.3, 5 pN, respectively.

**Figure 5A orange:** 4564, 4425, 4469, 4515, 4593, 4396, 4327, 4295 steps were used for 0.1, 0.32, 0.8, 1.5, 2.2, 2.9, 3.6, 4.3, 5 pN, respectively.

**Figure 5B red:** 1022, 1064, 888, 763, 720, 839, 784, 881 steps were used for 0.1, 0.2, 0.3, 0.4, 0.7, 1, 1.3, 1.6, 1.9, 2.2 pN, respectively.

**Figure 5B blue:** 140, 128, 136, 144, 151, 144, 133, 136 steps were used for 0.1, 0.2, 0.3, 0.4, 0.7, 1, 1.3, 1.6, 1.9, 2.2 pN, respectively.

**Figure 5B green:** 74, 74, 72, 82, 83, 78, 74, 71 steps were used for 0.1, 0.2, 0.3, 0.4, 0.7, 1, 1.3, 1.6, 1.9, 2.2 pN, respectively.

**Figure 5B orange:** 545, 534, 544, 583, 574, 601, 567, 565 steps were used for 0.1, 0.2, 0.3, 0.4, 0.7, 1, 1.3, 1.6, 1.9, 2.2 pN, respectively.

**Figure 5C red:** A total of 6813 steps were used. These were binned as: 1464, 1220, 1005, 1076, 1205, 737, 106 steps were used for relative extension 0.31, 0.4, 0.5, 0.6, 0.7, 0.8, 0.9, respectively.

**Figure 5C blue:** A total of 12951 steps were used. These were binned as: 4400, 3011, 2053, 1647, 1221, 545, 74 steps were used for relative extension 0.31, 0.41, 0.5, 0.6, 0.7, 0.79, 0.89, respectively.

**Figure 5C green:** A total of 11849 steps were used. These were binned as: 2804, 2449, 1820, 1804, 1820, 1035, 117 steps were used for relative extension 0.3, 0.4, 0.5, 0.61, 0.71, 0.81, 0.91, respectively.

**Figure 5C orange:** A total of 12014 steps were used. These were binned as: 3221, 2139, 1761, 1733, 1801, 1156, 203 steps were used for relative extension 0.32, 0.41, 0.51, 0.6, 0.69, 0.78, 0.88, respectively.

---

**Fig. S1.**
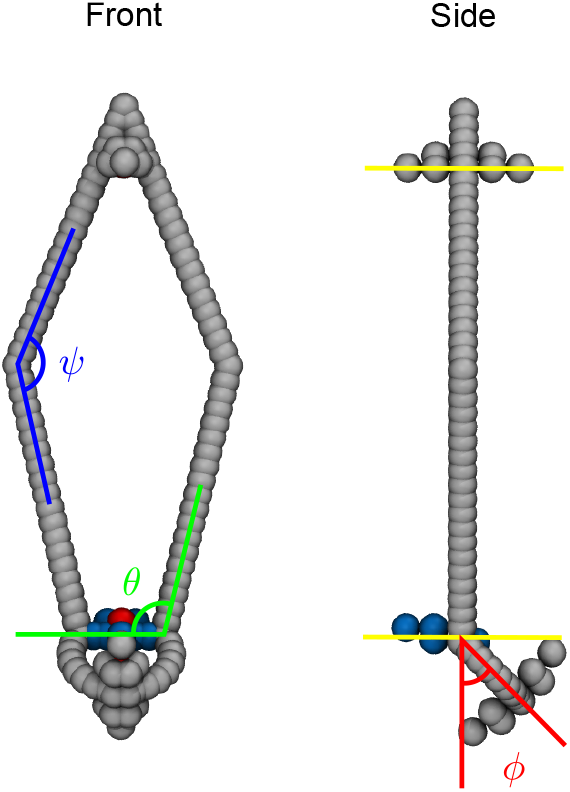
Interaction angles controlling the geometry of the SMCC (see Table S2). Additional dihedral interactions were used to fix the relative orientation of the top binding site and the ATP bridge (yellow lines). These have no physical meaning and are hence not shown.

**Fig. S2.**
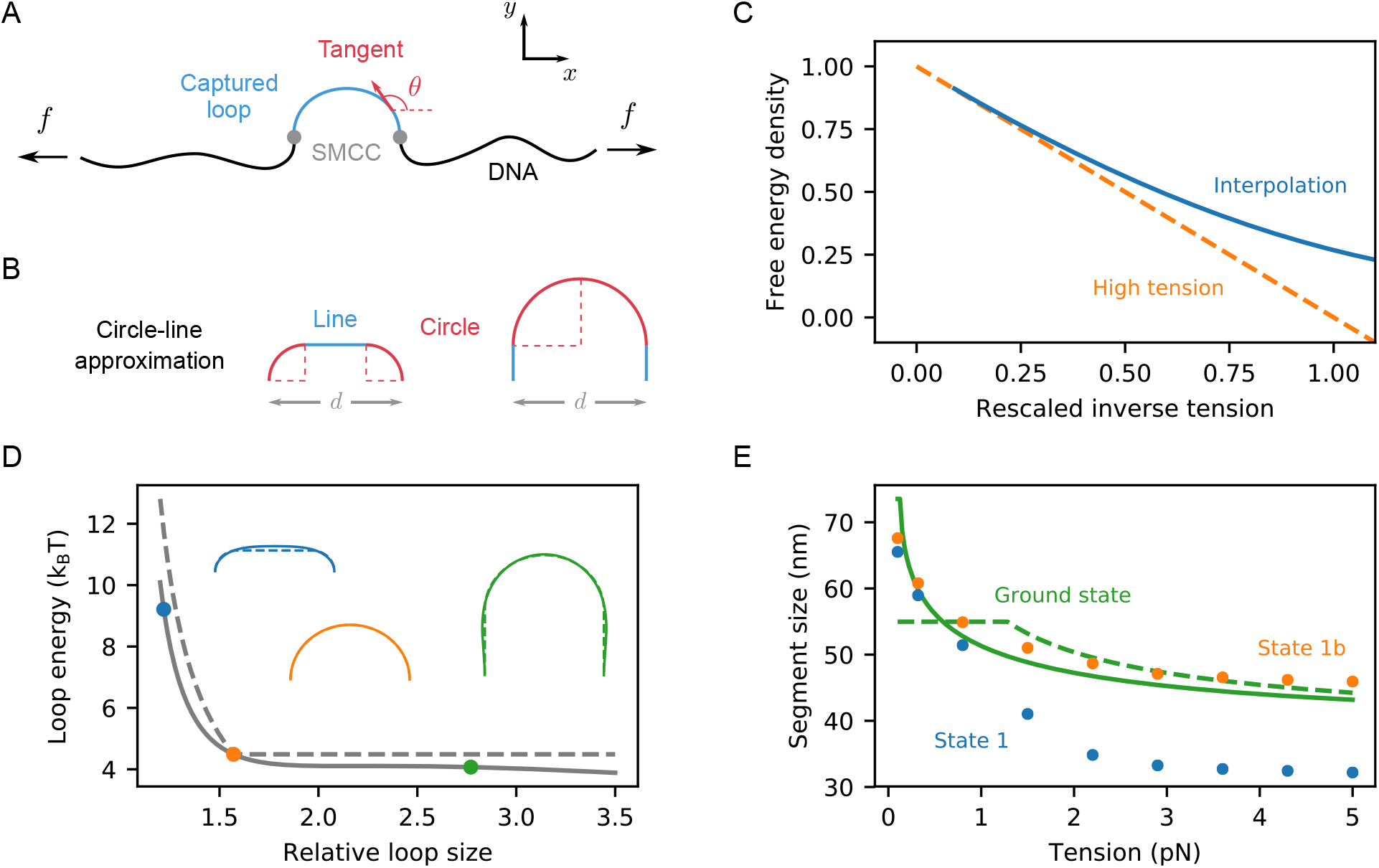
Ground-state calculation of captured segment size by SMCC. (A) During the ATP-bound state of the cycle, the SMCC model (shown with gray) captures a loop (blue line, see also state 1b in Fig. 1C of main text), the size of which depends on the applied DNA tension, *f* . Mathematically, the loop can be parametrized with a single angle, *θ*, which denotes the orientation of the tangent to the loop curve (indicated with red). (B) The captured loop can be described by a circle-line approximation, which consists of either two quadrants connected with a line (left) or a semicircle extended with two lines (right), depending on the loop length relative to its end-point distance, *d*. (C) Normalized free energy per unit length, −*g*(*f* )*/f* , of the stretched (*i.e.*, nonlooped) DNA, as a function of the dimensionless quantity (*Aβf* )^−1/2^. The solid, blue line is a numerical calculation, based on the interpolation formula of Ref. 11, while the dashed, orange line is high-force approximation, given by Eq. (2). (D) Ground-state energy, *ε*loop, *vs.* the relative loop size, *L/d*, together with the corresponding loop shapes for some selected values. The exact ground-state calculation [Eq. (7)] is shown with solid green line, while the circle-line approximation [Eq. (4)] with dashed green line. (E) Captured loop size *vs.* the applied DNA tension. Both the exact ground-state calculation (solid, green line) and the circle-line approximation (dashed, green line) seem to overestimate the simulation SMCC data (blue points) at high DNA tension. This deviation originates from partial-loop-capture events by the SMCC (state 1a in Fig. 1C of main text), which become increasingly probable at high tension. Excluding those from the calculation (orange points), reveals a good agreement with the ground-state theory.

**Fig. S3.**
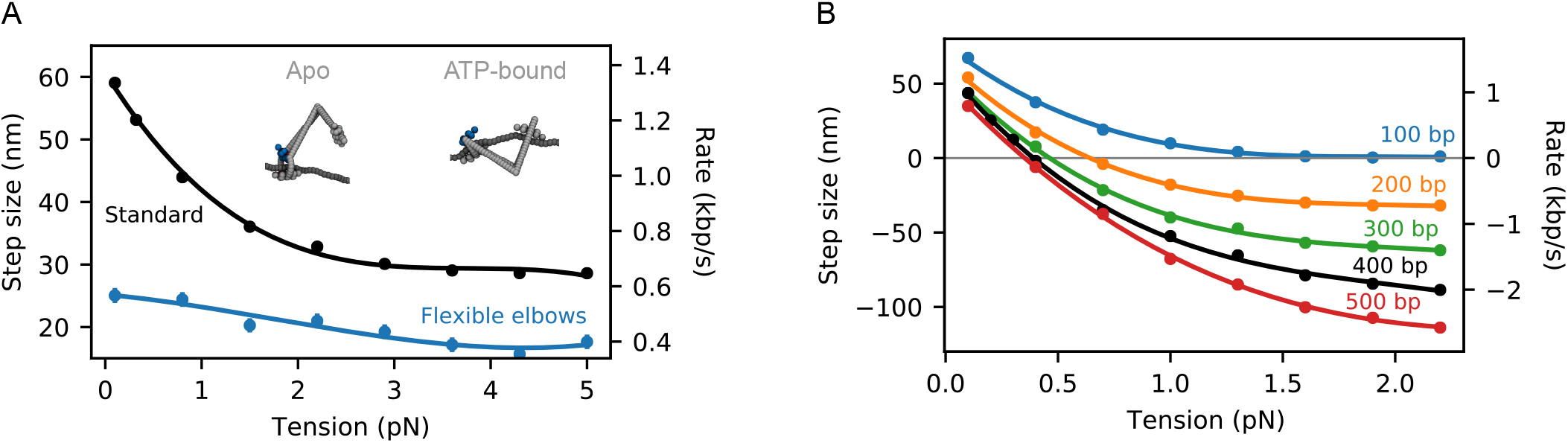
(A) Translocation step size *vs.* applied DNA tension, for completely flexible SMC arms. When the elbow stiffness is suppressed, translocation slows down (blue points), and exhibits a weak dependence on the DNA tension. Inspection of the simulated configurations (attached snapshots) reveals that, this is due to the SMCC folding in the ATP-bound state, rather than DNA looping. For comparison, the data of Fig. 2 of main text are also shown with black points, corresponding to semiflexible SMC elbows. (B) Dependence of the loop extrusion step size *vs.* applied DNA tension on the initial loop size. For comparison, the data of Fig. 3A of main text are shown with black points.

**Fig. S4.**
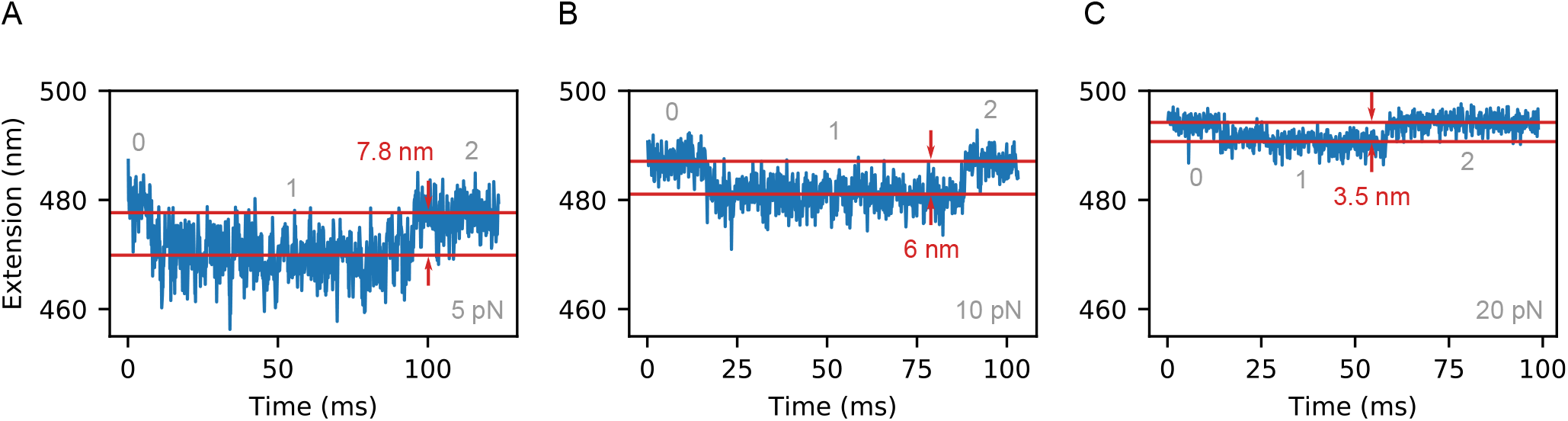
Time traces of DNA extension for a translocation cycle by SMCC, at DNA tension of (A) 5 pN, (B) 10 pN and (C) 20 pN. In all cases, the ATP-bound state (1) of the SMCC brings about a noticeable DNA contraction (indicated with red), which is a signature of the segment-capture mechanism (state 1 in Fig. 1C,D of main text). As expected, this is a decreasing function of DNA tension, as segment capture becomes increasingly unfavorable. In order to convert from simulation to real time units, we used a mean cycle duration of 0.13 s, as estimated in Fig. 3 of main text.

**Fig. S5.**
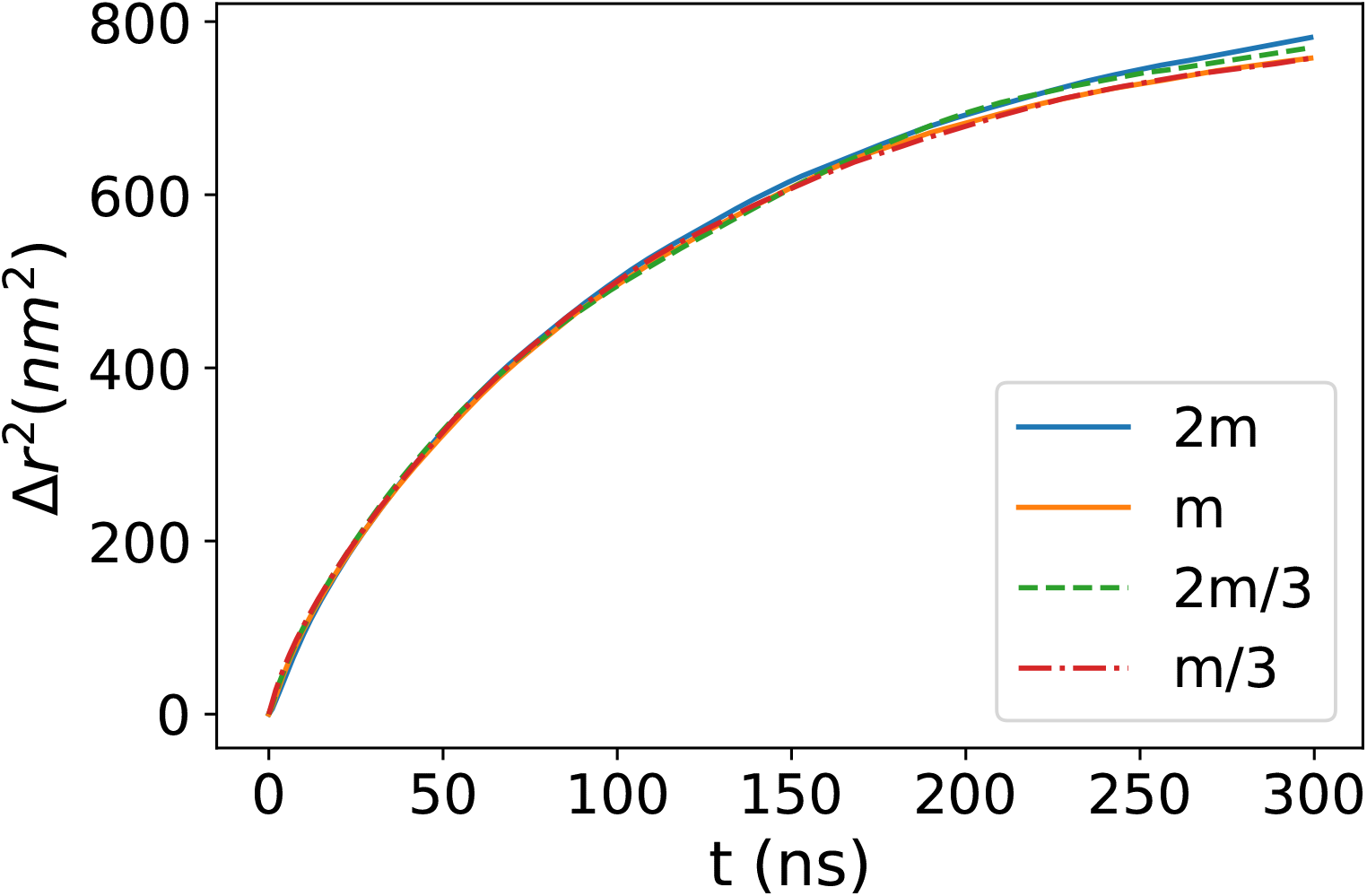
Mean squared displacement of the central monomer of a 400 bp (*L* = 136 nm) coarse-grained DNA loop vs time. The end-points of the loop are kept fixed at a distance of 23 nm. Simulations were repeated for different values of the mass of the coarse-grained beads *m* and damping coefficient *dm*, keeping the ratio *m/d_m_* constant. The data reported as *m* represent simulations with the parameters as used in the main text. The overlap of the data show that intertial effects are negligible beyond the time scale of 1 ns.

**Fig. S6.**
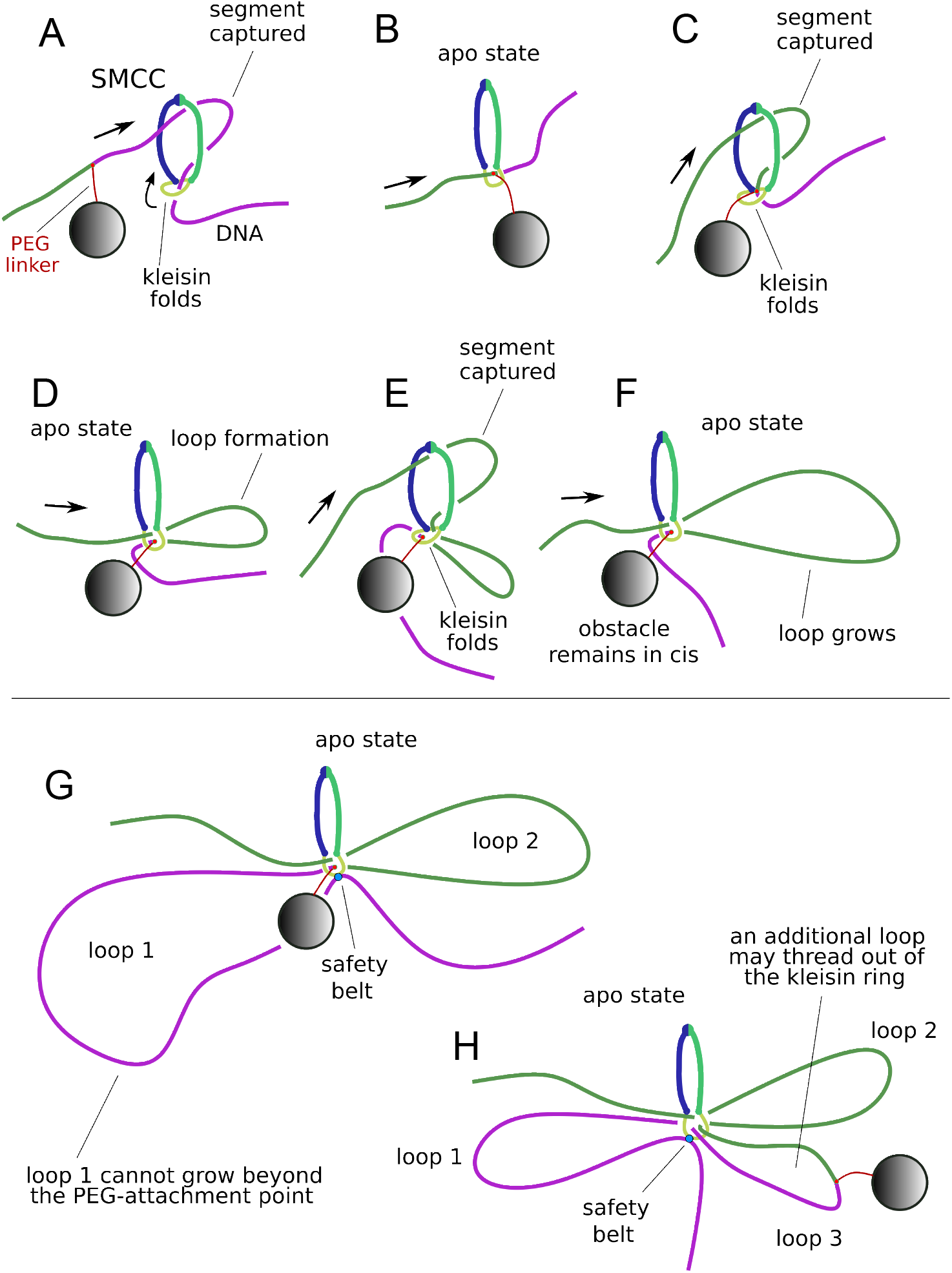
Schematic illustration of the interaction of a SMCC with a DNA to which a large nanoparticle is attached via a PEG-linker as in recent experiments (16). The figure illustrates how the SMCC handles the obstacle in the segment-capture model. For clarity we used different colors to identify different parts of the DNA. (A-F): The translocation process proceeds normally via segment-capture (A) until the obstacle reaches the basis of the SMCC (B). The obstacle remains then topologically trapped in the vicinity of the SMCC while the segment-capture mechanism can continue to operate (C) inducing the formation of a loop (D). As the process continues, captured segments (E) contribute to the growth of the loop (F). (G-H): Loop extrusion with external safety-belt mechanism (only the apo state is shown here). The extrusion proceeds regularly until the obstacle reaches the base of the SMCC. From that point on a secondary loop is extruded (G), as in translocation (D). The loop 1 cannot grow beyond the PEG-attachment point, but loop 2 keeps growing following the cycles of segment-capture as illustrated in (E) and (F). Note that the nanoparticle/obstacle need not remain in the vicinity of the SMCC (H), but another loop can thread out of the kleisin ring (loop 3). Also the same scheme applies to the cases where the anchoring point is outside the tripartite SMCC ring (topological loading) or where it is inside the ring (pseudotopological loading).

**Fig. S7.**
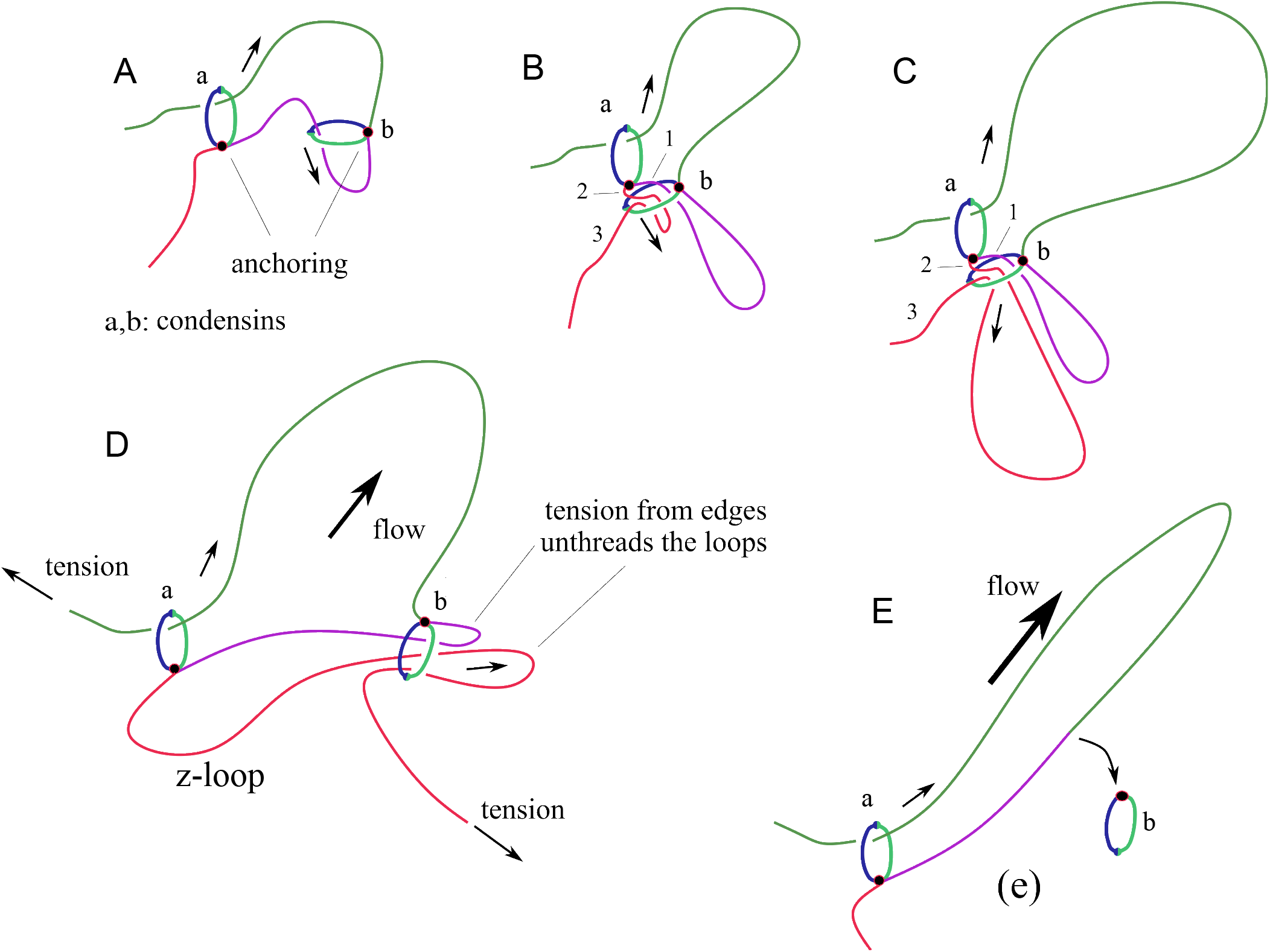
Condensins extruding loops on DNA were shown to be able to traverse one other (17). The figure illustrates a possible explanation of the apparent “crossing” based on the segment-capture model. (A) Two nested condensins (a and b) extruding loops asymmetrically, where the black dots denotes the anchoring point. The arrows show the direction of motion of the translocating strands. (B) While extruding the purple loop the condensin b approaches condensin a. This approach can lead to partial threading of a through b so that the red strand can be captured by condensin b. Three strands (shown as 1, 2 and 3) are in the ring. (C) The segment capture mechanism cannot work on strand 1 and 2 if these are too short, therefore condensin b pulls primarily strand 3, enlarging the red loop. (D) As the two endpoints of the DNA are attached to a solid surface in the experiments of Ref. (17), some tension builds up at the edges. This induces a separation of the two condensins from each other, generating the characteristic “Z-loop” shape observed under flow condition in the experiments (17). (E) The increasing tension and flow may lead to the unthreading of the loops from condensin b, leaving a single loop through a, as observed in experiments (17). This process can occur in either the case where the anchoring sites indicated are outside the tripartite SMCC ring (“safety belt” mechanism with topological loading, thought to be the case for yeast condensin) or inside the SMCC ring (“pseudo-topological loading).

**Movie S1. A typical translocation cycle for a DNA tension of 0.1 pN, showing the segment capture by the top binding site (state 1b)**.

**Movie S2. A typical translocation cycle for a DNA tension of 10 pN, showing the “inchworm”-like motion of the SMCC (state 1a).**

**Movie S3. Gradual loop extrusion of DNA with fixed end points by SMCC (10 cycles).**

## Notes

http://itf.fys.kuleuven.be/~enrico/SMC_SI/main.html

### Competing Interest Statement

The authors have declared no competing interest.

### Summary of Updates

Improved discussion of model and additional data. No substantial change in conclusions.

## REFERENCES

1. Nasmyth, K. (2001) Disseminating the genome: joining, resolving, and separating sister chromatids during mitosis and meiosis. Annu. Rev. Genet., 35(1), 673–745.

2. Alipour, E. and Marko, J. F. (2012) Self-organization of domain structures by DNA-loop-extruding enzymes. Nucleic Acids Res., 40(22), 11202–11212.

3. Datta, S., Lecomte, L., and Haering, C. H. (2020) Structural insights into DNA loop extrusion by SMC protein complexes. Curr. Opin. Struct. Biol., 65, 102–109.

4. Hirano, T. and Mitchison, T. J. (1994) A heterodimeric coiled-coil protein required for mitotic chromosome condensation in vitro. Cell, 79(3), 449–458.

5. Strunnikov, A. V., Hogan, E., and Koshland, D. (1995) SMC2, a Saccharomyces cerevisiae gene essential for chromosome segregation and condensation, defines a subgroup within the SMC family. Genes Dev., 9(5), 587–599.

6. Goloborodko, A., Marko, J. F., and Mirny, L. A. (2016) Chromosome compaction by active loop extrusion. Biophys. J., 110(10), 2162–2168.

7. Goloborodko, A., Imakaev, M. V., Marko, J. F., and Mirny, L. (2016) Compaction and segregation of sister chromatids via active loop extrusion. eLife, 5, e14864.

8. Gibcus, J. H., Samejima, K., Goloborodko, A., Samejima, I., Naumova, N., Nuebler, J., Kanemaki, M. T., Xie, L., Paulson, J. R., Earnshaw, W. C., Mirny, L. A., and Dekker, J. (2018) A pathway for mitotic chromosome formation. Science, 359(6376).

9. Strunnikov, A. V., Larionov, V. L., and Koshland, D. (1993) SMC1: an essential yeast gene encoding a putative head-rod-tail protein is required for nuclear division and defines a new ubiquitous protein family. J. Cell Biol., 123(6 Pt 2), 1635–1648.

10. Guacci, V., Koshland, D., and Strunnikov, A. (1997) A direct link between sister chromatid cohesion and chromosome condensation revealed through the analysis of MCD1 in S. cerevisiae. Cell, 91(1), 47–57.

11. Michaelis, C., Ciosk, R., and Nasmyth, K. (1997) Cohesins: chromosomal proteins that prevent premature separation of sister chromatids. Cell, 91(1), 35–45.

12. Sanborn, A. L., Rao, S. S., Huang, S.-C., Durand, N. C., Huntley, M. H., Jewett, A. I., Bochkov, I. D., Chinnappan, D., Cutkosky, A., Li, J., et al. (2015) Chromatin extrusion explains key features of loop and domain formation in wild-type and engineered genomes. Proc. Natl. Acad. Sci. U.S.A., 112(47), E6456–E6465.

13. Fudenberg, G., Imakaev, M., Lu, C., Goloborodko, A., Abdennur, N., and Mirny, L. A. (2016) Formation of chromosomal domains by loop extrusion. Cell Rep., 15(9), 2038–2049.

14. Hirano, M. and Hirano, T. (1998) ATP-dependent aggregation of single-stranded DNA by a bacterial SMC homodimer. EMBO J., 17(23), 7139–7148.

15. Wang, X., Brandão, H. B., Le, T. B., Laub, M. T., and Rudner, D. Z. (2017) Bacillus subtilis SMC complexes juxtapose chromosome arms as they travel from origin to terminus. Science, 355(6324), 524–527.

16. Niki, H., Jaffe, A., Imamura, R., Ogura, T., and Hiraga, S. (1991) The new gene mukB codes for a 177 kd protein with coiled-coil domains involved in chromosome partitioning of E. coli.. EMBO J., 10(1), 183–193.

17. Lioy, V. S., Cournac, A., Marbouty, M., Duigou, S., Mozziconacci, J., Espéli, O., Boccard, F., and Koszul, R. (2018) Multiscale structuring of the E. coli chromosome by nucleoid-associated and condensin proteins. Cell, 172(4), 771–783.

18. Mäkelä, J. and Sherratt, D. J. (2020) Organization of the Escherichia coli chromosome by a MukBEF axial core. Mol. Cell, 78, 250–260.

19. Diebold-Durand, M.-L., Lee, H., Avila, L. B. R., Noh, H., Shin, H.-C., Im, H., Bock, F. P., Bürmann, F., Durand, A., Basfeld, A., et al. (2017) Structure of full-length SMC and rearrangements required for chromosome organization. Mol. Cell, 67(2), 334–347.

20. Lee, B. G., Merkel, F., Allegretti, M., Hassler, M., Cawood, C., Lecomte, L., O’Reilly, F. J., Sinn, L. R., Gutierrez-Escribano, P., Kschonsak, M., Bravo, S., Nakane, T., Rappsilber, J., Aragon, L., Beck, M., L?we, J., and Haering, C. H. (2020) Cryo-EM structures of holo condensin reveal a subunit flip-flop mechanism. Nat Struct Mol Biol, 27(8), 743–751.

21. Hirano, T. (2016) Condensin-Based Chromosome Organization from Bacteria to Vertebrates. Cell, 164(5), 847–857.

22. Haering, C. H., Farcas, A. M., Arumugam, P., Metson, J., and Nasmyth, K. (2008) The cohesin ring concatenates sister DNA molecules. Nature, 454(7202), 297–301.

23. Cuylen, S., Metz, J., and Haering, C. H. (2011) Condensin structures chromosomal DNA through topological links. Nat. Struct. Mol. Biol., 18(8), 894–901.

24. Wilhelm, L., Bürmann, F., Minnen, A., Shin, H. C., Toseland, C. P., Oh, B. H., and Gruber, S. (2015) SMC condensin entraps chromosomal DNA by an ATP hydrolysis dependent loading mechanism in Bacillus subtilis. Elife, 4.

25. Nunez, R. V., Avila, L. B. R., and Gruber, S. (2019) Transient DNA occupancy of the SMC interarm space in prokaryotic condensin. Mol. Cell, 75(2), 209–223.

26. Shi, Z., Gao, H., Bai, X. C., and Yu, H. (2020) Cryo-EM structure of the human cohesin-NIPBL-DNA complex. Science, 368(6498), 1454–1459.

27. Bürmann, F., Lee, B.-G., Than, T., Sinn, L., O’Reilly, F. J., Yatskevich, S., Rappsilber, J., Hu, B., Nasmyth, K., and Löwe, J. (2019) A folded conformation of MukBEF and cohesin. Nat. Struct. Mol. Biol., 26(3), 227–236.

28. Bürmann, F., Funke, L. F. H., Chin, J. W., and Löwe, J. (2021) Cryo-EM structure of MukBEF reveals DNA loop entrapment at chromosomal unloading sites. Mol. Cell, 81(23), 4891–4906.

29. Serrano, D., Cordero, G., Kawamura, R., Sverzhinsky, A., Sarker, M., Roy, S., Malo, C., Pascal, J. M., Marko, J. F., and D’Amours, D. (2020) The Smc5/6 Core Complex Is a Structure-Specific DNA Binding and Compacting Machine. Mol. Cell, 80(6), 1025–1038.

30. Soh, Y. M., Bürmann, F., Shin, H. C., Oda, T., Jin, K. S., Toseland, C. P., Kim, C., Lee, H., Kim, S. J., Kong, M. S., Durand-Diebold, M. L., Kim, Y. G., Kim, H. M., Lee, N. K., Sato, M., Oh, B. H., and Gruber, S. (2015) Molecular basis for SMC rod formation and its dissolution upon DNA binding. Mol. Cell, 57(2), 290–303.

31. Kschonsak, M., Merkel, F., Bisht, S., Metz, J., Rybin, V., Hassler, M., and Haering, C. H. (2017) Structural basis for a safety-belt mechanism that anchors condensin to chromosomes. Cell, 171(3), 588–600.

32. Strick, T. R., Kawaguchi, T., and Hirano, T. (2004) Real-time detection of single-molecule DNA compaction by condensin I. Curr. Biol., 14(10), 874–880.

33. Eeftens, J. M., Bisht, S., Kerssemakers, J., Kschonsak, M., Haering, C. H., and Dekker, C. (2017) Real-time detection of condensin-driven DNA compaction reveals a multistep binding mechanism. EMBO J., 36(23), 3448–3457.

34. Keenholtz, R. A., Dhanaraman, T., Palou, R., Yu, J., D’Amours, D., and Marko, J. F. (2017) Oligomerization and ATP stimulate condensin-mediated DNA compaction. Sci. Rep., 7(1), 14279.

35. Terakawa, T., Bisht, S., Eeftens, J. M., Dekker, C., Haering, C. H., and Greene, E. C. (2017) The condensin complex is a mechanochemical motor that translocates along DNA. Science, 358(6363), 672–676.

36. Ganji, M., Shaltiel, I. A., Bisht, S., Kim, E., Kalichava, A., Haering, C. H., and Dekker, C. (2018) Real-time imaging of DNA loop extrusion by condensin. Science, 360(6384), 102–105.

37. Davidson, I. F., Bauer, B., Goetz, D., Tang, W., Wutz, G., and Peters, J. M. (2019) DNA loop extrusion by human cohesin. Science, 366(6471), 1338–1345.

38. Kim, Y., Shi, Z., Zhang, H., Finkelstein, I. J., and Yu, H. (2019) Human cohesin compacts DNA by loop extrusion. Science, 366(6471), 1345–1349.

39. Golfier, S., Quail, T., Kimura, H., and Brugués, J. (2020) Cohesin and condensin extrude DNA loops in a cell cycle-dependent manner. Elife, 9.

40. Kong, M., Cutts, E. E., Pan, D., Beuron, F., Kaliyappan, T., Xue, C., Morris, E. P., Musacchio, A., Vannini, A., and Greene, E. C. (2020) Human Condensin I and II Drive Extensive ATP-Dependent Compaction of Nucleosome-Bound DNA. Mol. Cell, 79(1), 99–114.

41. Marko, J. F., De Los Rios, P., Barducci, A., and Gruber, S. (2019) DNA-segment-capture model for loop extrusion by structural maintenance of chromosome (SMC) protein complexes. Nucleic Acids Res., 47(13), 6956–6972.

42. Nichols, M. H. and Corces, V. G. (2018) A tethered-inchworm model of SMC DNA translocation. Nat. Struct. Mol. Biol., 25(10), 906–910.

43. Lawrimore, J., Friedman, B., Doshi, A., and Bloom, K. (2017) RotoStep: a chromosome dynamics simulator reveals mechanisms of loop extrusion. In Cold Spring Harb. Symp. Quant. Biol. Cold Spring Harbor Laboratory Press Vol. 82, pp. 101–109.

44. Brackley, C. A., Johnson, J., Michieletto, D., Morozov, A. N., Nicodemi, M., Cook, P. R., and Marenduzzo, D. (Sep, 2017) Nonequilibrium Chromosome Looping via Molecular Slip Links. Phys. Rev. Lett., 119, 138101.

45. Bonato, A. and Michieletto, D. (2021) Three-dimensional loop extrusion. Biophys. J, 120(24), 5544–5552.

46. Takaki, R., Dey, A., Shi, G., and Thirumalai, D. (2021) Theory and simulations of condensin mediated loop extrusion in DNA. Nat Commun, 12(1), 5865.

47. Ryu, J.-K., Katan, A. J., van der Sluis, E. O., Wisse, T., de Groot, R., Haering, C. H., and Dekker, C. (2020) The condensin holocomplex cycles dynamically between open and collapsed states. Nat. Struct. Mol. Biol., pp. 1–8.

48. Hassler, M., Shaltiel, I. A., and Haering, C. H. (2018) Towards a Unified Model of SMC Complex Function. Curr Biol, 28(21), R1266–R1281.

49. Bauer, B. W., Davidson, I. F., Canena, D., Wutz, G., Tang, W., Litos, G., Horn, S., Hinterdorfer, P., and Peters, J. M. (2021) Cohesin mediates DNA loop extrusion by a ”swing and clamp” mechanism. Cell, 184(21), 5448–5464.

50. Yu, Y., Li, S., Ser, Z., Sanyal, T., Choi, K., Wan, B., Kuang, H., Sali, A., Kentsis, A., Patel, D. J., and Zhao, X. (2021) Integrative analysis reveals unique structural and functional features of the Smc5/6 complex. Proc Natl Acad Sci U S A, 118(19).

51. Taschner, M., Basquin, J., Steigenberger, B., Schäfer, I. B., Soh, Y. M., Basquin, C., Lorentzen, E., Räschle, M., Scheltema, R. A., and Gruber, S. (2021) Nse5/6 inhibits the Smc5/6 ATPase and modulates DNA substrate binding. EMBO J, 40(15), e107807.

52. Petela, N. J., Gonzalez Llamazares, A., Dixon, S., Hu, B., Lee, B. G., Metson, J., Seo, H., Ferrer-Harding, A., Voulgaris, M., Gligoris, T., Collier, J., Oh, B. H., Löwe, J., and Nasmyth, K. A. (2021) Folding of cohesin’s coiled coil is important for Scc2/4-induced association with chromosomes. Elife, 10.

53. Ryu, J. K., Rah, S.-H., Janissen, R., Kerssemkers, J. W. J., and Dekker, C. (2020) Resolving the step size in condensin driven DNA loop-extrusion identifies ATP binding as the step-generating process. biorXiv, p. 2020.11.04.368506v1.

54. Hirano, M. and Hirano, T. (2002) Hinge-mediated dimerization of SMC protein is essential for its dynamic interaction with DNA. EMBO J., 21(21), 5733–5744.

55. Chiu, A., Revenkova, E., and Jessberger, R. (2004) DNA interaction and dimerization of eukaryotic SMC hinge domains. J. Biol. Chem., 279(25), 26233–26242.

56. Griese, J. J., Witte, G., and Hopfner, K. P. (2010) Structure and DNA binding activity of the mouse condensin hinge domain highlight common and diverse features of SMC proteins. Nucleic Acids Res., 38(10), 3454–3465.

57. Kurze, A., Michie, K. A., Dixon, S. E., Mishra, A., Itoh, T., Khalid, S., Strmecki, L., Shirahige, K., Haering, C. H., Löwe, J., and Nasmyth, K. (2011) A positively charged channel within the Smc1/Smc3 hinge required for sister chromatid cohesion. EMBO J, 30(2), 364–378.

58. Sun, M., Nishino, T., and Marko, J. F. (2013) The SMC1-SMC3 cohesin heterodimer structures DNA through supercoiling-dependent loop formation. Nucleic Acids Res., 41(12), 6149–6160.

59. Uchiyama, S., Kawahara, K., Hosokawa, Y., Fukakusa, S., Oki, H., Nakamura, S., Kojima, Y., Noda, M., Takino, R., Miyahara, Y., Maruno, T., Kobayashi, Y., Ohkubo, T., and Fukui, K. (2015) Structural Basis for Dimer Formation of Human Condensin Structural Maintenance of Chromosome Proteins and Its Implications for Single-stranded DNA Recognition. J. Biol. Chem., 290(49), 29461–29477.

60. Alt, A., Dang, H. Q., Wells, O. S., Polo, L. M., Smith, M. A., McGregor, G. A., Welte, T., Lehmann, A. R., Pearl, L. H., Murray, J. M., and Oliver, A. W. (2017) Specialized interfaces of Smc5/6 control hinge stability and DNA association. Nat. Commun., 8, 14011.

61. Chapard, C., Jones, R., van Oepen, T., Scheinost, J. C., and Nasmyth, K. (2019) Sister DNA entrapment between juxtaposed smc heads and kleisin of the cohesin complex. Mol. Cell, 75(2), 224–237.

62. Collier, J. E., Lee, B. G., Roig, M. B., Yatskevich, S., Petela, N. J., Metson, J., Voulgaris, M., Gonzalez Llamazares, A., Löwe, J., and Nasmyth, K. A. (2020) Transport of DNA within cohesin involves clamping on top of engaged heads by Scc2 and entrapment within the ring by Scc3. Elife, 9.

63. Seifert, F. U., Lammens, K., Stoehr, G., Kessler, B., and Hopfner, K.-P. (2016) Structural mechanism of ATP-dependent DNA binding and DNA end bridging by eukaryotic Rad50. EMBO J., 35(7), 759–772.

64. Liu, Y., Sung, S., Kim, Y., Li, F., Gwon, G., Jo, A., Kim, A.-K., Kim, T., Song, O.-k., Lee, S. E., et al. (2016) ATP-dependent DNA binding, unwinding, and resection by the Mre11/Rad50 complex. EMBO J., 35(7), 743–758.

65. Higashi, T. L., Eickhoff, P., Sousa, J. S., Locke, J., Nans, A., Flynn, H. R., Snijders, A. P., Papageorgiou, G., O’Reilly, N., Chen, Z. A., O’Reilly, F. J., Rappsilber, J., Costa, A., and Uhlmann, F. (2020) A Structure-Based Mechanism for DNA Entry into the Cohesin Ring. Mol. Cell, 79(6), 917–933.

66. Shaltiel, I. A., Datta, S., Lecomte, L., Hassler, M., Kschonsak, M., Bravo, S., Stober, C., Eustermann, S., and Haering, C. H. (2021) A hold- and-feed mechanism drives directional DNA loop extrusion by condensin. bioRxiv,.

67. Gligoris, T. G., Scheinost, J. C., Bürmann, F., Petela, N., Chan, K. L., Uluocak, P., Beckouèt, F., Gruber, S., Nasmyth, K., and Löwe, J. (2014) Closing the cohesin ring: structure and function of its Smc3-kleisin interface. Science, 346(6212), 963–967.

68. Banigan, E. J., van den Berg, A. A., Brandõ, H. B., Marko, J. F., and Mirny, L. A. (2020) Chromosome organization by one-sided and two-sided loop extrusion. Elife, 9.

69. Muir, K. W., Li, Y., Weis, F., and Panne, D. (2020) The structure of the cohesin ATPase elucidates the mechanism of SMC-kleisin ring opening. Nat. Struct. Mol. Biol., 27(3), 233–239.

70. Plimpton, S. (1995) Fast parallel algorithms for short-range molecular dynamics. J. Comput. Phys., 117(1), 1–19.

71. Rybenkov, V. V., Cozzarelli, N. R., and Vologodskii, A. V. (1993) Probability of DNA knotting and the effective diameter of the DNA double helix. Proc. Natl. Acad. Sci. U.S.A., 90(11), 5307–5311.

72. Bouchiat, C., Wang, M. D., Allemand, J.-F., Strick, T., Block, S., and Croquette, V. (1999) Estimating the persistence length of a worm-like chain molecule from force-extension measurements. Biophys. J., 76(1), 409–413.

73. He, Y., Lawrimore, J., Cook, D., Van Gorder, E. E., De Larimat, S. C., Adalsteinsson, D., Forest, M. G., and Bloom, K. (2020) Statistical mechanics of chromosomes: in vivo and in silico approaches reveal high-level organization and structure arise exclusively through mechanical feedback between loop extruders and chromatin substrate properties. Nucl. Acids Res., 48(20), 11284–11303.

74. Stigler, J., Camdere, G. Ö., Koshland, D. E., and Greene, E. C. (2016) Single-molecule imaging reveals a collapsed conformational state for DNA-bound cohesin. Cell Rep., 15(5), 988–998.

75. Brandão, H. B., Paul, P., van den Berg, A. A., Rudner, D. Z., Wang, X., and Mirny, L. A. (2019) RNA polymerases as moving barriers to condensin loop extrusion. Proc. Natl. Acad. Sci. U.S.A., 116(41), 20489–20499.

76. Eeftens, J. M., Katan, A. J., Kschonsak, M., Hassler, M., de Wilde, L., Dief, E. M., Haering, C. H., and Dekker, C. (2016) Condensin Smc2-Smc4 Dimers Are Flexible and Dynamic. Cell Rep., 14(8), 1813–1818.

77. Giuntoli, R. D., Linzer, N. B., Banigan, E. J., Sing, C. E., De La Cruz, M. O., Graham, J. S., Johnson, R. C., and Marko, J. F. (2015) DNA-segment-facilitated dissociation of Fis and NHP6A from DNA detected via single-molecule mechanical response. J. Mol. Biol., 427(19), 3123–3136.

78. Badrinarayanan, A., Reyes-Lamothe, R., Uphoff, S., Leake, M. C., and Sherratt, D. J. (2012) In vivo architecture and action of bacterial structural maintenance of chromosome proteins. Science, 338(6106), 528–531.

79. Zawadzka, K., Zawadzki, P., Baker, R., Rajasekar, K. V., Wagner, F., Sherratt, D. J., and Arciszewska, L. K. (2018) MukB ATPases are regulated independently by the N- and C-terminal domains of MukF kleisin. Elife, 7.

80. Rajasekar, K. V., Baker, R., Fisher, G. L. M., Bolla, J. R., Mäkelä, J., Tang, M., Zawadzka, K., Koczy, O., Wagner, F., Robinson, C. V., Arciszewska, L. K., and Sherratt, D. J. (2019) Dynamic architecture of the Escherichia coli structural maintenance of chromosomes (SMC) complex, MukBEF. Nucleic Acids Res., 47(18), 9696–9707.

81. Pradhan, B., Barth, R., Kim, E., Davidson, I. F., Bauer, B., van Laar, T., Yang, W., Ryu, J.-K., van der Torre, J., Peters, J.-M., and Dekker, C. (2021) SMC complexes can traverse physical roadblocks bigger than their ring size. bioRxiv,.

82. Higashi, T. L. and Uhlmann, F. (2022) SMC complexes: Lifting the lid on loop extrusion. Curr Opin Cell Biol, 74, 13–22.

83. Kim, E., Kerssemakers, J., Shaltiel, I. A., Haering, C. H., and Dekker, C. (2020) DNA-loop extruding condensin complexes can traverse one another. Nature, 579(7799), 438–442.

84. Brandão, H. B., Ren, Z., Karaboja, X., Mirny, L. A., and Wang, X. (2021) DNA-loop-extruding SMC complexes can traverse one another in vivo.

## References

1. M Hirano, T Hirano, Opening closed arms: long-distance activation of SMC ATPase by hinge-DNA interactions. Mol Cell 21, 175–186 (2006).

2. RV Nunez, LBR Avila, S Gruber, Transient DNA occupancy of the SMC interarm space in prokaryotic condensin. Mol. Cell 75, 209–223 (2019).

3. JJ Griese, G Witte, KP Hopfner, Structure and DNA binding activity of the mouse condensin hinge domain highlight common and diverse features of SMC proteins. Nucleic Acids Res. 38, 3454–3465 (2010).

4. S Uchiyama, et al., Structural Basis for Dimer Formation of Human Condensin Structural Maintenance of Chromosome Proteins and Its Implications for Single-stranded DNA Recognition. J. Biol. Chem. 290, 29461–29477 (2015).

5. S Datta, L Lecomte, CH Haering, Structural insights into DNA loop extrusion by SMC protein complexes. Curr. Opin. Struct. Biol. 65, 102–109 (2020).

6. H Koide, N Kodera, S Bisht, S Takada, T Terakawa, Modeling of DNA binding to the condensin hinge domain using molecular dynamics simulations guided by atomic force microscopy. bioRxiv (2021).

7. Y Liu, et al., ATP-dependent DNA binding, unwinding, and resection by the Mre11/Rad50 complex. EMBO J. 35, 743–758 (2016).

8. Y Li, et al., Structural basis for Scc3-dependent cohesin recruitment to chromatin. eLife 7, e38356 (2018).

9. M Kschonsak, et al., Structural basis for a safety-belt mechanism that anchors condensin to chromosomes. Cell 171, 588–600 (2017).

10. JF Marko, Biophysics of protein–DNA interactions and chromosome organization. Phys. A 418, 126–153 (2015).

11. C Bouchiat, et al., Estimating the persistence length of a worm-like chain molecule from force-extension measurements. Biophys. J. 76, 409–413 (1999).

12. I Kulić, H Schiessel, Nucleosome repositioning via loop formation. Biophys. J. 84, 3197–3211 (2003).

13. S Sankararaman, JF Marko, Formation of loops in DNA under tension. Phys. Rev. E 71, 021911 (2005).

14. SK Nomidis, et al., Twist-bend coupling, twist waves, and the shape of DNA loops. Phys. Rev. E 100, 022402 (2019).

15. S Plimpton, Fast parallel algorithms for short-range molecular dynamics. J. Comput. Phys. 117, 1–19 (1995).

16. B Pradhan, et al., Smc complexes can traverse physical roadblocks bigger than their ring size. bioRxiv (2021).

17. E Kim, J Kerssemakers, IA Shaltiel, CH Haering, C Dekker, DNA-loop extruding condensin complexes can traverse one another. Nature 579, 438–442 (2020).

